# Drug-dependent morphological transitions in spherical and worm-like polymeric micelles define stability and pharmacological performance of micellar drugs

**DOI:** 10.1101/2021.06.10.447962

**Authors:** Chaemin Lim, Jacob D. Ramsey, Duhyeong Hwang, Susana C. M. Teixeira, Chi-Duen Poon, Joshua D Strauss, Marina Sokolsky-Papkov, Alexander V. Kabanov

## Abstract

Significant advances in physicochemical properties of polymeric micelles enable optimization of therapeutic drug efficacy, supporting nanomedicine manufacturing and clinical translation. Yet, the effect of micelle morphology on pharmacological efficacy has not been adequately addressed. We addressed this gap by assessing pharmacological efficacy of polymeric micelles with spherical and worm-like morphologies. We observed that poly(2-oxazoline)-based polymeric micelles can be elongated over time from a spherical structure to worm-like structure, with elongation influenced by several conditions, including the amount and type of drug loaded into the micelles. We further evaluated the role of different morphologies of olaparib micelles on pharmacological performance against a triple-negative breast cancer tumor (TNBC) model. Spherical micelles accumulated rapidly in the tumor tissue while retaining large amounts of drug; worm-like micelles accumulated more slowly and only upon releasing significant amounts of drug. These findings suggest that the dynamic character of the drug–micelle structure and the micelle morphology play a critical role in pharmacological performance, and that spherical micelles are better suited for systemic delivery of anticancer drugs to tumors when drugs are loosely associated with the polymeric micelles.

## 1. Introduction

Polymeric micelles formed by amphiphilic block copolymers exhibit large potential as platforms for solubilization and delivery of poorly soluble therapeutic compounds. Several polymeric-micelle drugs are approved for clinical use and others are in the pipeline for clinical evaluation and approval worldwide. Unique traits of polymeric micelles include ease of self-assembly, universality with respect to drug compatibility, and ability to address major challenges in pharmaceutics, such as poor drug solubility. However, micelles’ dynamic character also limits *in vivo* testing and therefore understanding of their performance. Despite considerable progress in this field, understanding of how the physicochemical properties (stability, size, shape, extent of drug loading, and drug release) ^[1-3]^ relate to the pharmacological performance remains elusive.

Kataoka and colleagues demonstrated that tumor accumulation of polymeric micelles depends on particle size and tumor desmoplasticity ^[2]^; smaller polymeric micelles show better accumulation and penetration into more desmoplastic tumor and possess improved antitumor activity. A study by Discher and colleagues emphasized the importance of micelle morphology; filament, worm-like micelles display greater longevity in plasma circulation, whereas micelle disassembly into smaller spherical particles results in faster clearance ^[1]^. Yet, neither of these studies related micellar pharmacological performance to their drug loading and release characteristics—a critical aspect of drug efficacy. In the former study, drug was covalently bound to the micelles and slowly released as a result of ligand-exchange, a process proceeding in biological milieu over several days. In the latter, micelles were free of the drug. Subsequent attempts to prepare micelles that maintain filament morphology while retaining pharmacologically relevant drug loadings have been unsuccessful ^[4]^.

A useful and potentially universal embodiment of this technology is the polymeric micelle system that may (i) solubilize drugs without the need for chemical drug modification, (ii) have a drug-loading capacity of at least 10 % by mass, and (iii) release drug within at least twelve hours (i.e., slow release). The current study advances our understanding of the pharmacological performance of such micelles by using poly(2-oxazoline) (POx) block copolymers. These block copolymers form polymeric micelles that incorporate many structurally diverse drugs at high loadings. The differential, non-covalent interactions of the block copolymers with the drugs leads to the formation of micelles with two distinct morphological and stabilization characteristics: (i) those that stabilize small spherical micelles, and (ii) those that allow the formation of worm-like micelles, even at high drug loadings. We also examined the complex interplay between micelle circulation, drug release, and the ability of different morphologies to reach their site of action while carrying the loaded drug. To do so, we carried out a comprehensive analysis that includes physiochemical structural measurements, molecular interactions and modeling, drug release, *in vivo* pharmacokinetics (PK), and anti-tumor activity in animal models. Our findings indicate that formulating drugs into small polymeric micelles is beneficial to their pharmacological performance; their geometries are conducive to increased colloidal stability, formulation reproducibility, and drug delivery to tumors.

## 2. Results

### 2.1 Multiple morphologies in blank polymeric micelles

To understand how micellar morphologies influence drug-loading and pharmacological performance, we prepared micellar structures with varying morphologies. This was accomplished by self-assembly of amphiphilic block copolymers above a critical micelle concentration (CMC). Micellar structures contain a hydrophobic core that is shielded from the ambient aqueous solution by a corona of hydrophilic blocks. Micellar structures can exhibit different shapes, including spheres, rods, worm-like structures, and bilayers which can subsequently develop into vesicles ^[5, 6]^. In particular, polymeric micelles can undergo one-dimensional (1-D) growth to form long worm-like aggregates. This behavior is captured in the four synthesized A-B-A triblock copolymers (T1–T4; **Supplementary Figure S1**), in which hydrophilic poly(2-methyl-2-oxazoline) A-blocks vary in length and hydrophobic poly(2-butyl-2-oxazoline) B-blocks are kept constant at 20 repeating units (r.u.). Micelles were prepared from these copolymers using a thin film hydration method; resulting structures were observed using transmission electron microscopy (TEM). We also examined them at two time intervals: 1 hour and 72 hours after formation. After 1 hour, all micelles exhibited small spherical structures **(Figure 1a)**. After 72 hours, the population gradually shifted to worm-like morphologies **(Figure 1b** and **Supplementary Figure S2a**). Further, the elongation process from spherical to worm-like micelles strongly correlated to the change in particle size and size distribution, as shown by dynamic light scattering (DLS; **Figure 1c**). Spherical particles examined after 1 hour exhibited particle sizes in the 10–30 nm range with a narrow size distribution. As micelles elongated, particle size and PDI value increased simultaneously, indicating the co-existence of spherical micelles and worm-like micelles. Micelles continued to elongate until reaching worm-like structures 200 nm or larger and exhibiting a narrow size distribution. Thus, DLS provides a simple tool for predicting the morphology of nanoparticles. Generally, elongation was either favored by increasing the polymer concentration and temperature, or inhibited by increasing the ionic strength (see **Supplementary Figure S2b**). Elongation and growth were predominantly dependent on the A block length; the T1 and T2 polymers completely converted to non-spherical or worm-like micelles after 72 hours, whereas T3 and T4 transitioned to a mixture of morphologies. These results are consistent with the well-known principles for polymeric micelle formation ^[5, 7]^. The repulsion of the hydrophilic chains strongly influences the growth and the resulting morphology of the polymeric micelles; an increase in the hydrophilic block length led to an increase of the repulson due to the excluded volume effect within the shell, and to partial inhibition of the spherical-to-worm-like structural transitions in T3 and T4. Despite being thermodynamically favored, the worm-like formation process was kinetically limited, taking up to a couple of days before completion in T1 and T2.

**Figure 1.**
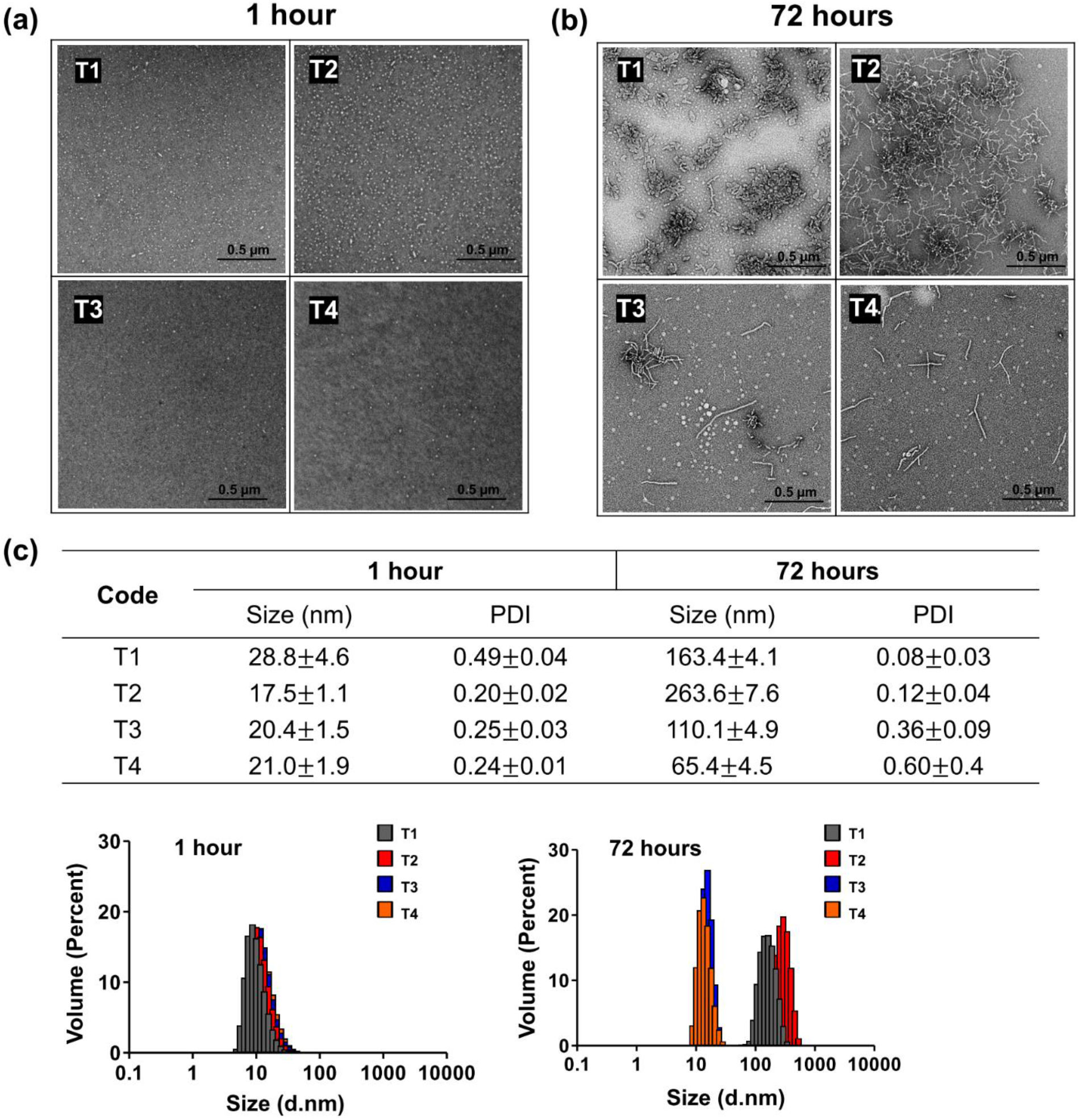
**(a, b)** TEM images and **(c)** DLS characteristics (Size (D_eff_, diameter), PDI, volume weighted size distribution) of T1–T4 based micelles at 1 hour and 72 hours. Polymer concentrations are 10 mg/mL in distilled water. Errors shown represent three standard deviations of uncertainty. (D_eff_ = the effective diffusion coefficient, PDI = Polydispersity index)

### 2.2. Drug dependent effects on micelle morphology

Previously, we showed that POx micelles can exhibit different morphologies when loaded with different drugs. We observed two types of behavior: the first, exemplified by paclitaxel, inhibited the formation of worm-like micelle architecture and stabilized the compact micelle spheres to retain a narrow size distribution ^[4]^; the second, exemplified by etoposide, was unable to prevent the formation of elongated worm-like micelles even at high drug loadings ^[8]^. For simplicity, we refer henceforth to the first behavior as “worm-inhibiting”, and to the second behavior as “worm-permissive”. As a proof of concept, we randomly selected a set of 20 poorly soluble drugs and tested their loading efficiency (LE) and loading capacity (LC) in our POx micelles **(Supplementary Table S1)**. Drugs with high LC were first encapsulated by POx micelles at various loading ratios (polymer/drug w/w ratios of 10/1, 10/2, 10/4 and 10/8). Then, the sphere-to-worm-like transition processes were monitored over time as done before drug-loading. For comparison we also varied the temperature at 4°C and 25° C, which strongly affected the transition, and used the T2 block copolymer with relatively shorter hydrophilic blocks to increase the propensity of the transitions **(Figure 2**).

**Figure 2.**
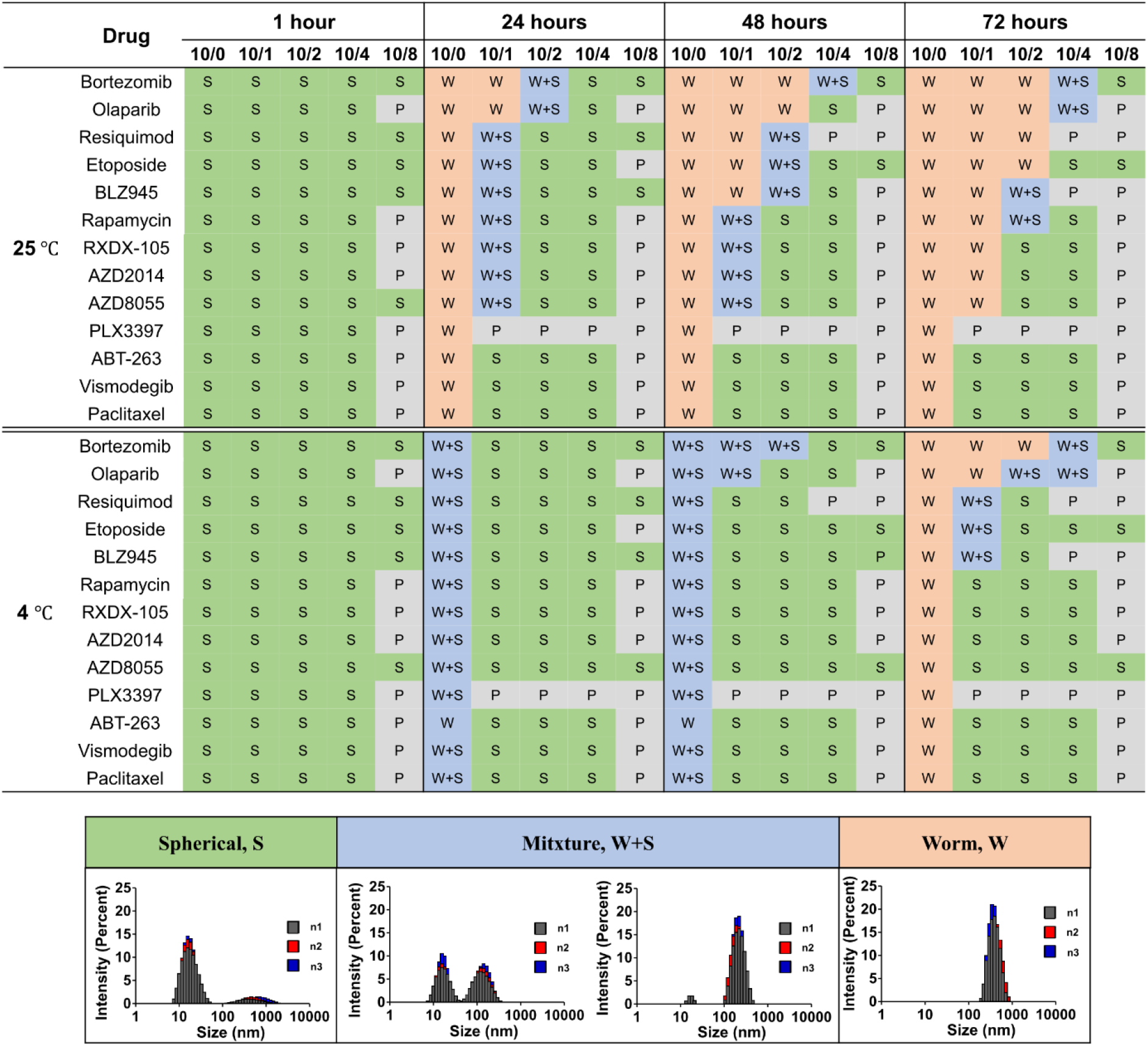
Monitoring the sphere-to-worm-like transition process of T2-based polymeric micelles at various drug loading ratios (polymer/drug w/w ratios of 10/1, 10/2, 10/4 and 10/8) over time. The matrix denotes the resulting morphologies: S, spherical micelles; W+S, mixture of worms and spheres; W, worm-like micelles; and P, precipitation. Increased drug loading resulted in slowing or inhibition of the sphere-to-worm transition.

We next examined the role of size and synthesis conditions on micellar structure. DLS was used as an indicator of resulting morphologies, which were then validated by cryo-TEM (**Supplementary Figures S3** and **S4**). Generally, sphere-to-worm transitions were slowed or inhibited as drug loading increased. This was evidenced by the reduced width of peaks in the distribution of hydrodynamic diameters. Drugs exhibited differential ability to inhibit worm formation. For example, at room temperature, blank spherical micelles become elongated into worm structures after a 24-hour incubation. [The sphere-to-worm transition rarely happened at low temperature (4 °C).] Micelles loaded with bortezomib, olaparib, etoposide, or resiquimod showed a similar or slightly slower transition to worm-like structure, which was the most pronounced at 72 hours. Other drugs, such as BLZ945, rapamycin, RXDX105, AZD2014, and AZD8055, slightly inhibited the sphere-to-worm transition (especially at higher loading) but did not abolish the transition. In contrast, vismodegib, paclitaxel, and ABT-263 inhibited elongation at a mass fraction of as little as 10 %. Therefore, we unveiled two distinct groups of drugs: those that are worm-permissive under certain conditions (even at high drug loadings) and those that are worm-inhibiting and stabilize the compact spherical micelle morphology. Notably for most of these drugs, the formation of worms and in many cases precipitates at high loadings were suppressed by using a block copolymer with longer hydrophilic blocks, such as T3 (**Supplementary Figure S5**). Interestingly, for the worm-inhibiting drugs ABT-263, paclitaxel, and vismodegib, maximal solubilization in T3 was two-fold more than that of the same drugs in T2 (8 mg/mL vs. 4 mg/mL). Solubility of the worm-permissive drugs did not differ for these polymers.

### 2.3. Analysis of micelle structure by SANS

To better understand the morphology and effects of different drugs in POx micelles, small-angle neutron scattering (SANS) data were collected on a series of drug concentrations while keeping the block copolymer concentration constant (10 mg/mL T2 in D_2_O). **Figure 3** shows the scattering data and fits at different drug loadings. A polydisperse core-shell spherical model for the single micelles in solution provides a consistently reasonable fit for the medium to high scattering angle data. Both vismodegib and paclitaxel stabilize core-shell spherical particles at lower drug concentrations, while worm-like larger clusters still contribute to scattering in the presence of olaparib and etoposide up to 2 mg/mL of drug.

**Figure 3.**
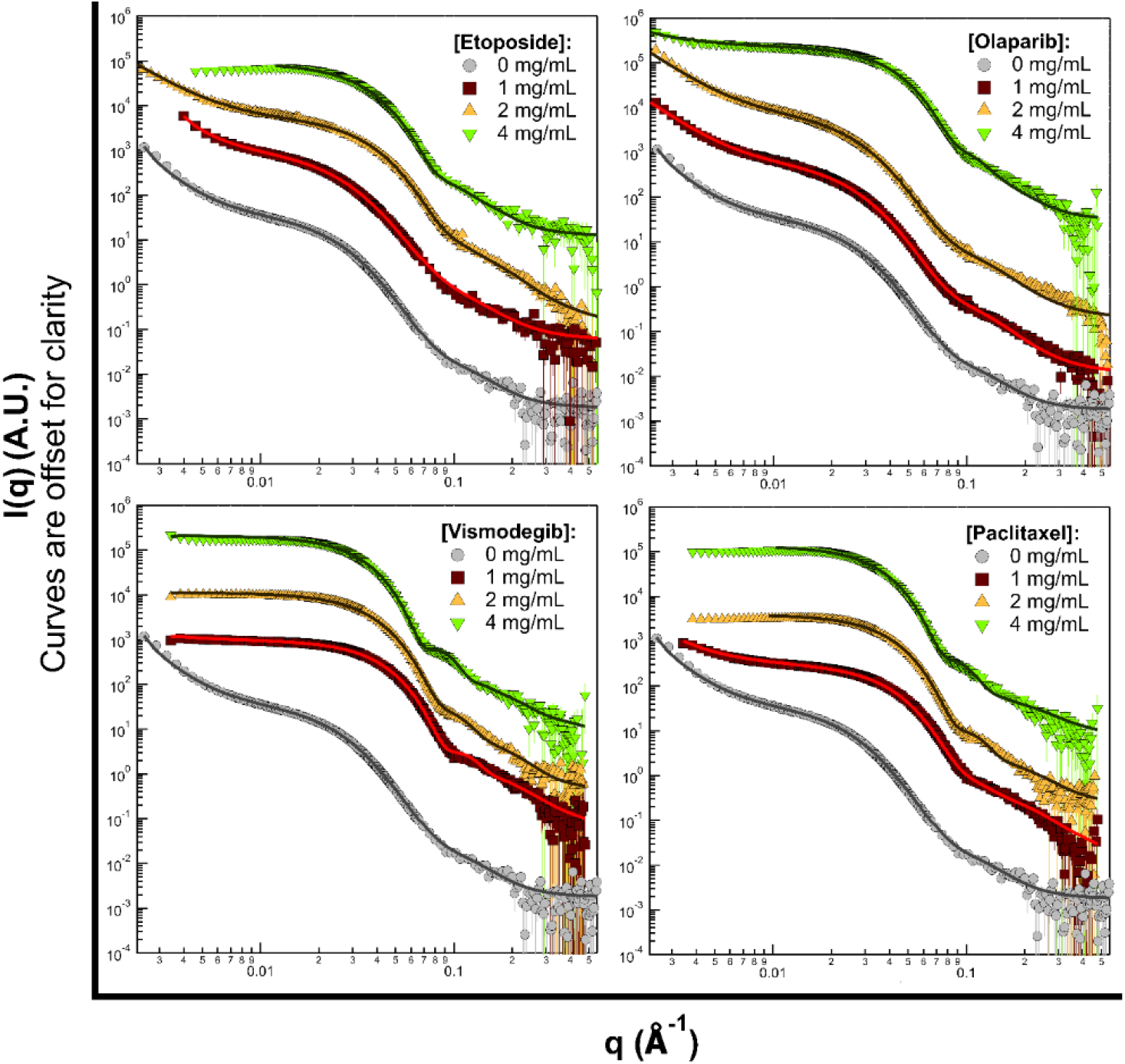
SANS data (symbols) measured for solutions of 10 mg/mL of T2 with various drug loading ratios, as labeled. The continuous lines show fits to the data, where the model function was smeared with the instrument resolution information prior to fitting. Error bars correspond to one standard deviation of uncertainty.

A comparative plot of the average core radius is shown in **Figure 4**, shell thickness and neutron scattering length densities (SLD) of micelles, as a function of the drug concentration (see **Supplementary Table S2** for a list of the fit parameters). Although the data fitting can only provide a semi-quantitative measurement, given the polydispersity of the samples and the presence of larger particles with dimensions outside the scope of the SANS data measured, the results are consistent with our findings for the worm-permissive and worm-inhibiting drugs. The calculated average diameter of blank T2 sphere micelles in solution is 14.2 ± 1.1 nm (where 1.1 nm corresponds to one standard deviation of uncertainty). With increasing drug concentration, all T2 solutions show a growth in the core radius, with no significant variation in the core SLD. When compared to the worm-permissive drugs, paclitaxel and vismodegib show more pronounced core size increases and smaller shell thickness. The observed shell SLD with drug loading is consistent with increased hydration of the shell. For the worm-permissive drugs, the shell SLD appears to increase proportionally to drug concentration. Overall, the variations observed in core size, shell thickness and SLDs reflect rearrangements of the micelle composition and density in the presence of increasing amounts of drug.

**Figure 4.**
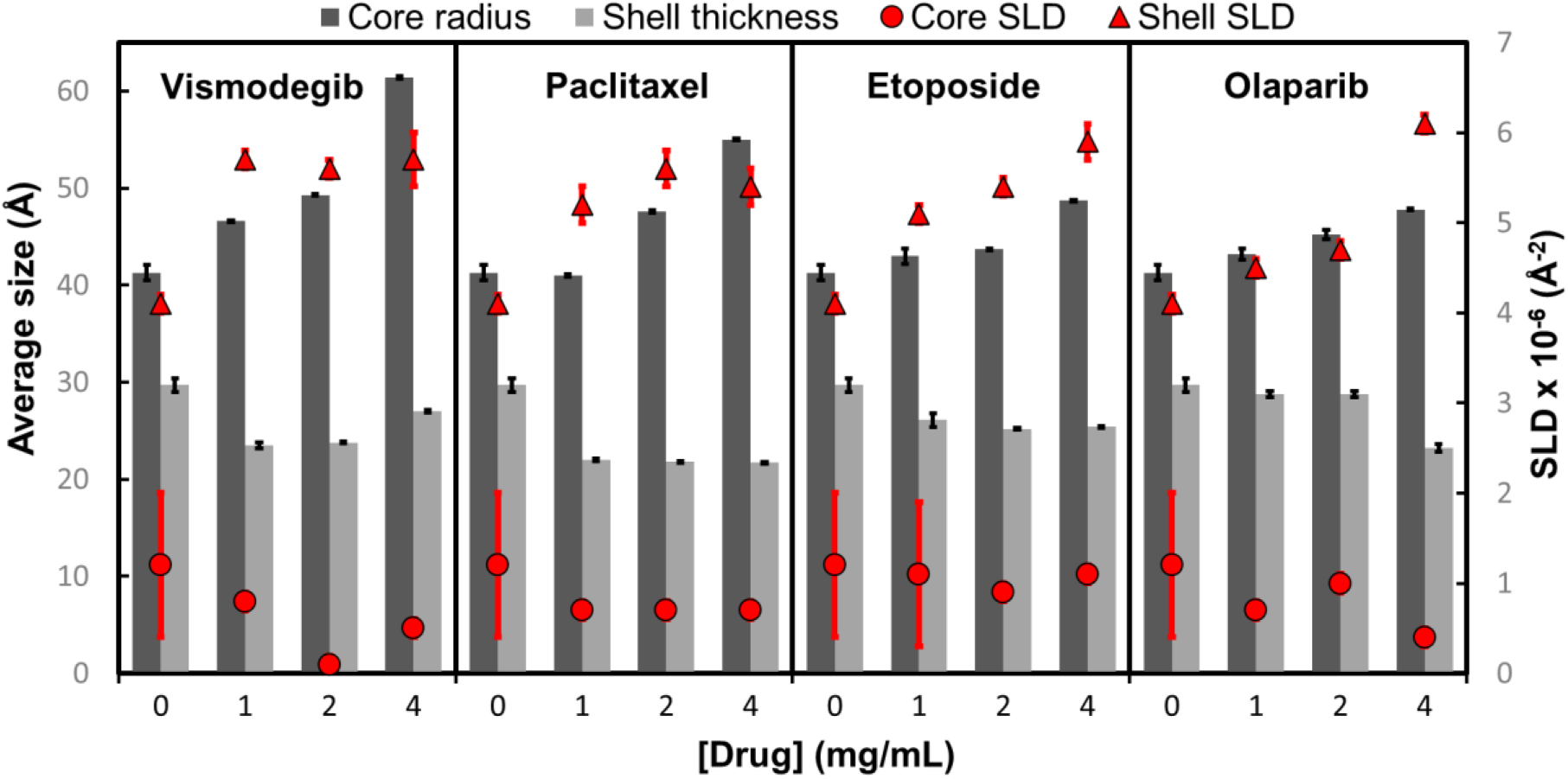
Fitted parameters for SANS data of T2 micelles in the presence of various drug concentrations. The dark grey bars represent the average core radius of the micelles in the sample, the light grey bars show the average shell thickness, red circles show the micelle core SLD, and red squares show the shell SLD. For comparison, the calculated neutron SLD for pure drug solutions in 100 % D_2_O are: 2.929 × 10^−6^ Å ^-2^ for Etoposide (d = 1.6 g/cm^3^), 2.797 × 10^−6^ Å ^-2^ for olaparib (d = 1.4 g/cm^3^), 2.694 × 10^−6^ Å ^-2^ for paclitaxel (d = 1.4 g/cm^3^), and 2.845 × 10^−6^ Å ^-2^ for vismodegib (d = 1.4 g/cm^3^). Error bars correspond to one standard deviation of uncertainty.

### 2.4. Assessing drug localization in the micelles by MD simulation

To understand drug locations within the micellar structures, we performed molecular dynamics (MD) simulations of drug-loaded T2 micellar structures. Simulations of etoposide and paclitaxel loaded into POx micelles were performed at various polymer and drug molar ratios (50:50, 50:100, and 50:200) corresponding to the mass ratios (10/1, 10/2, and 10/4) used throughout this work **(Figure 5)**. We observed self-aggregation of polymers in every simulation. At every drug loading, most etoposide molecules were located at the center of the micelles, indicating that etoposide binds to the hydrophobic micelle core. In contrast, paclitaxel was located initially in the peripheral areas of micelles at lower loading, potentially coming in contact with both core-and shell-forming blocks. As drug loading increased, drug placement in the core also increased. This simulation suggests large differences in the drug-polymer interactions displayed by worm-permitting and worm-inhibiting drugs.

**Figure 5.**
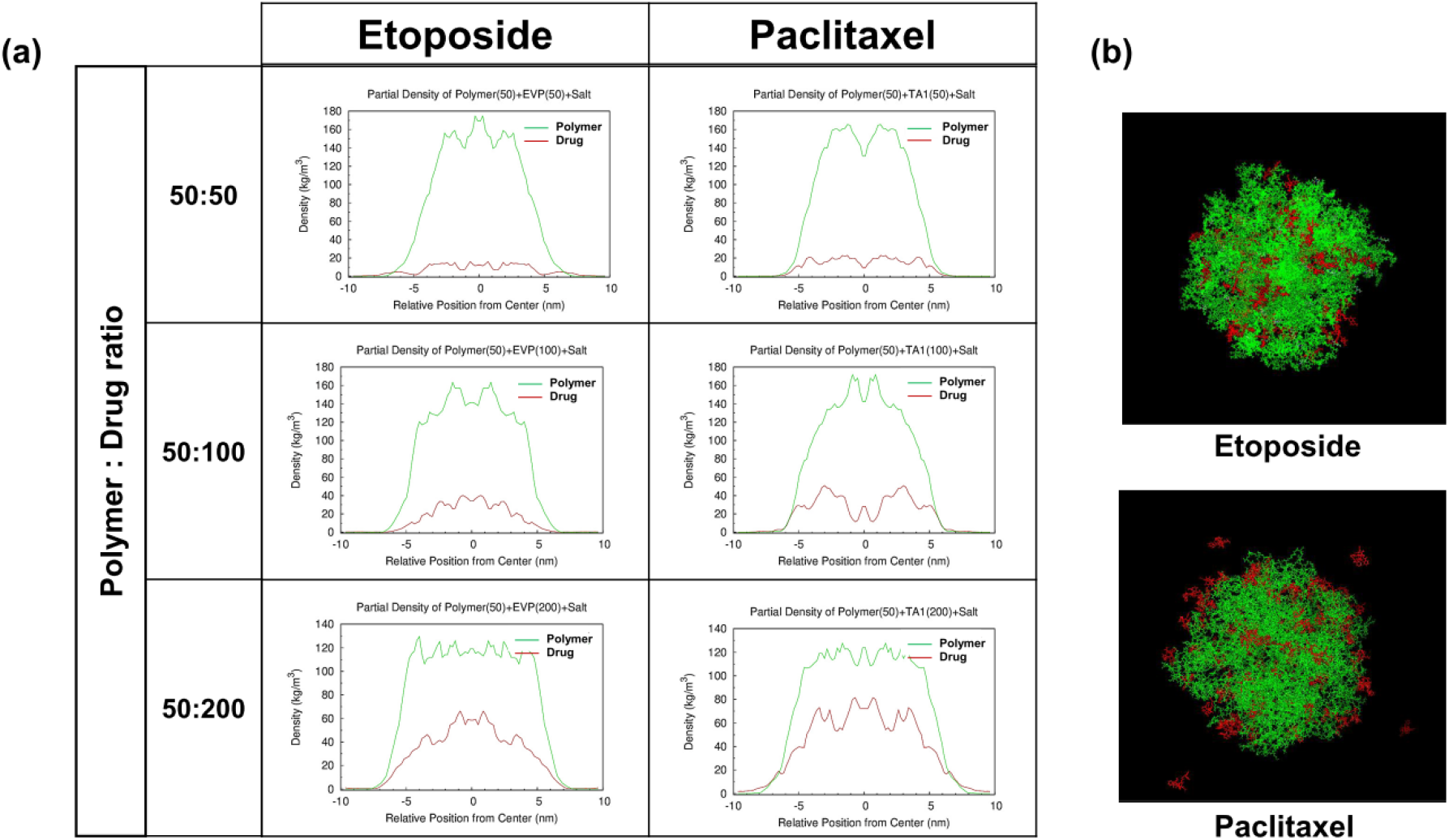
MD simulation of etoposide and paclitaxel in T2 micelles: **(a)** the drug (red) and polymer (green) density profiles from the center of the micelles, and **(b)** simulated snapshots of the drug-loaded micelles. The fifty-polymer chains and 50, 100, and 200 drug molecules were simulated for a total of 100 ns.

### 2.5. Effect of drugs on the stability of the micelles by DLS measurements

For the first time, we demonstrate here the differential effects of drugs on the stability of morphologically distinct micelles formed from the same block copolymer. We determined the effective CMC* by measuring DLS count rates upon dilution at 1 h and 72 h (see **Supplementary Figure S6**). The CMC* of the blank worm-like micelles was approximately six times smaller than that of spherical micelles for the same polymer (T2) (**Figure 6**).

**Figure 6.**
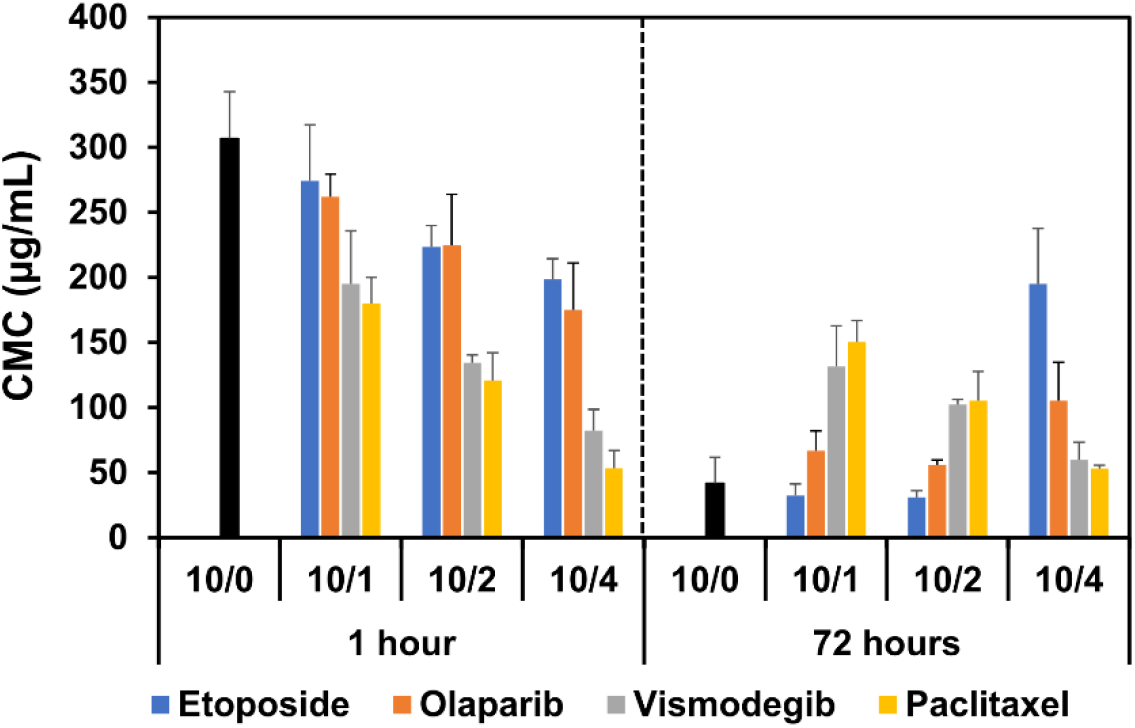
Effective CMC* of polymeric micelles at various drug loading ratios (polymer/drug ratios of 10/0, 10/1, 10/2, and 10/4) determined by DLS at 1 hour and 72 hours after formation at 25°C. Error bars correspond to three standard deviation of uncertainty.

The observed difference in effective CMC* suggests that the worm-like micelles of T2 are more stable than the spherical micelles in the absence of drugs. For worm-permissive olaparib and etoposide, we measured the effective CMC* for both morphologies at the same drug loading. Although the drug loading resulted in a decrease in CMC* for spherical micelles, worm-like micelles generally exhibited a much lower CMC* (**Figure 6**). Thus, worm-like micelles loaded with worm-permissive drugs appeared to be more stable than their spherical counterparts. In contrast, worm-inhibiting drugs (paclitaxel and vismodegib) decreased the CMC* of spherical micelles to a greater extent. At high drug loadings, the CMC* of spherical micelles both at 1 h and 72 h was as low as that of the worm-like micelles in the absence of drug. Thus, worm-inhibiting drugs appear to stabilize the spherical micelle structure upon dilution.

### 2.6. Probing drug effects on the micelle microenvironment by Reichardt’s dye

To investigate the changes of the micellar microenvironment upon the loading of different drugs we used 2,6-diphenyl-4-(2,4,6-triphenyl-N-pyridino)phenolate, better known as Reichardt’s dye (RD), which exhibits strong solvatochromatic effects in its absorbance spectrum ^[9]^. Generally, the RD incorporation in the blank, drug-free micelles resulted in a decrease in polarity due to the relative hydrophobic environment maintained by the 2-butyl-2-oxazoline core. This manifested in the absorbance spectrum as an increase in the dye λ_max_ compared to its absorbance in solution with polar solvents (see **Supplementary Figure S7a**,**b**). As the worm-inhibiting drugs paclitaxel and vismodegib were solubilized in the micelles, the dye environment became more polar, which led to profound decreases of the λ_max_ even at low drug loadings. In contrast, the worm-permissive drugs bortezomib, marimastat, resiquimod, and olaparib had little or no effect on the polarity, as evidenced by the lack of or slight changes in the λ_max_. Interestingly, etoposide (worm-permissive) had an intermediate effect, causing relatively smaller λ_max_ changes at low loading and changes comparable to those exhibited by paclitaxel and vismodegib at high loadings (**Figure 7**). Drugs also had strikingly different effects on the absorbance intensity of the RD, which characterizes the intermolecular interactions between the dye and surrounding molecules. The absorbance intensity of the dye decreased in the absence of drug upon solubilization in the micelles, likely due to hydrogen bonding and ion-dipole interactions of the dye with the polymer repeating units (**Supplementary Figure S7c**). Worm-permissive drugs either did not change the absorbance intensity (like olaparib, resiquimod, and etoposide) or further decrease it (like bortezomib and marimastat) (**Figure 7**).

**Figure 7.**
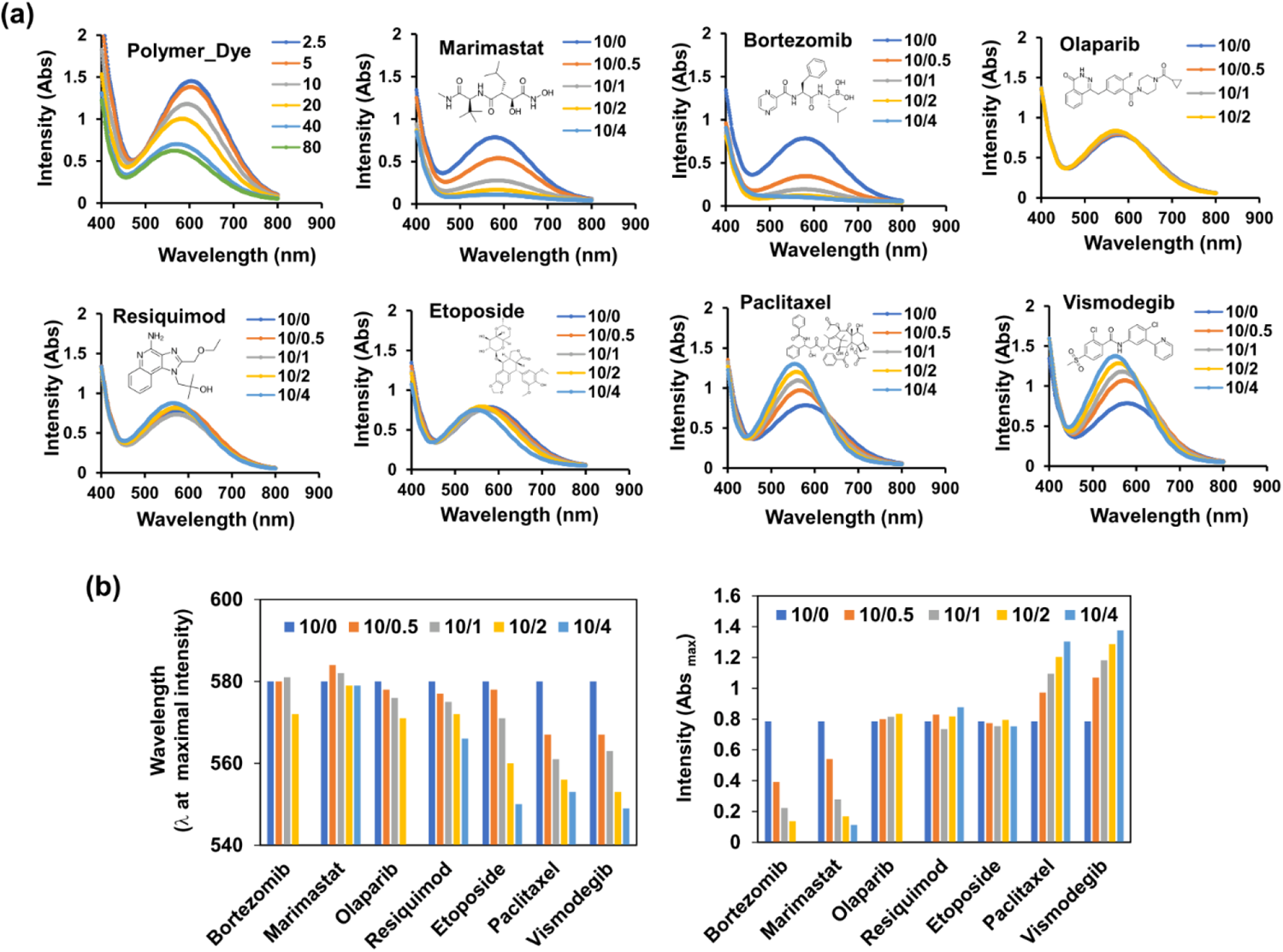
Characterization of micelle microenvironment by Reichardt’s dye (RD). **(a)** UV-Vis absorption spectra of RD in blank micelles (polymer/RD ratios of 2.5/1, 5/1, 10/1, 20/1, 40/1, and 80/1) and drug-loaded micelles (polymer/dye/drug ratios of 10/0.5/0, 10/0.5/1, 10/0.5/2, and 10/0.5/4). **(b)** Wavelength and intensity of RD obtained from UV-Vis absorption spectra.

This behavior potentially results from differential molecular interactions of RD with the polymer and drugs. The betaine-containing RD can participate in ion-dipole and hydrogen bonding interactions with adjacent non-ionic molecules containing peptide-bond motifs ^[10]^, and can likely engage in similar interactions with the poly(2-oxazoline) repeating units. Consequently, the absorbance intensity of RD is suppressed upon its incorporation in the micelles. When marimastat and bortezomib containing repeated amide bonds are added, the absorbance intensity further decreases—likely a result of direct interactions with RD. Other worm-permissive drugs such as etoposide, resiquimod, and olaparib lacking the amide bonds have no effect on the absorbance intensity of RD. Worm-inhibiting drugs paclitaxel and vismodegib contain numerous electronegative and hydrogen bond-accepting groups that may strengthen their interaction with the poly(2-oxazoline). Notably, NMR spectra previously suggested that paclitaxel can interact with the amide bond motif of both poly(2-methyl-2-oxasoline) and poly(2-butyl-2-oxasoline) ^[11]^. Therefore, such drugs can bind to the same binding sites that RD binds to and may result in the displacement of the dye if their binding affinity exceeds that of RD. This displacement of RD with the drugs would explain the observed increase in the absorbance intensity (see **Supplementary Figure S8**).

### 2.7. Preparing drug-loaded spherical and worm-like micelles for injections

Using this knowledge, we were able to alter the morphology of polymeric micelles while keeping drug loading constant, enabling us to thoroughly evaluate the effect of morphology on drug pharmacological performance. We selected olaparib, a poly(ADP-ribose) polymerase (PARP) inhibitor approved for treatment of advanced ovarian and metastatic breast cancer, as a model drug and formulated it into T2 polymeric micelles. Micelles were incubated at room temperature and monitored in a similar fashion to all other micelles to record any changes in particle size and morphology (**Figure 8a**,**b**). Over time, olaparib micelles elongated until forming worm-like structures by 72 hours; there were no changes in drug concentration or loading during elongation **(Figure 8c)**. Then, we examined whether changes in morphology affected the drug-release profiles in the micelles—a critical parameter for drug delivery. Notably, different morphologies of olaparib-loaded micelles exhibited identical *in vitro* drug release profiles **(Figure 8d)**. However, size and shape differences resulted in varying micellar mobility in a diffusion-constrained environment (e.g., agarose gel). A model experiment illustrated the improved penetration of such spherical micelles higher viscosity systems (see **Supplementary Figure S9**). Based on this, we proceeded with evaluating olaparib formulations of micelles with different morphologies in an animal model.

**Figure 8.**
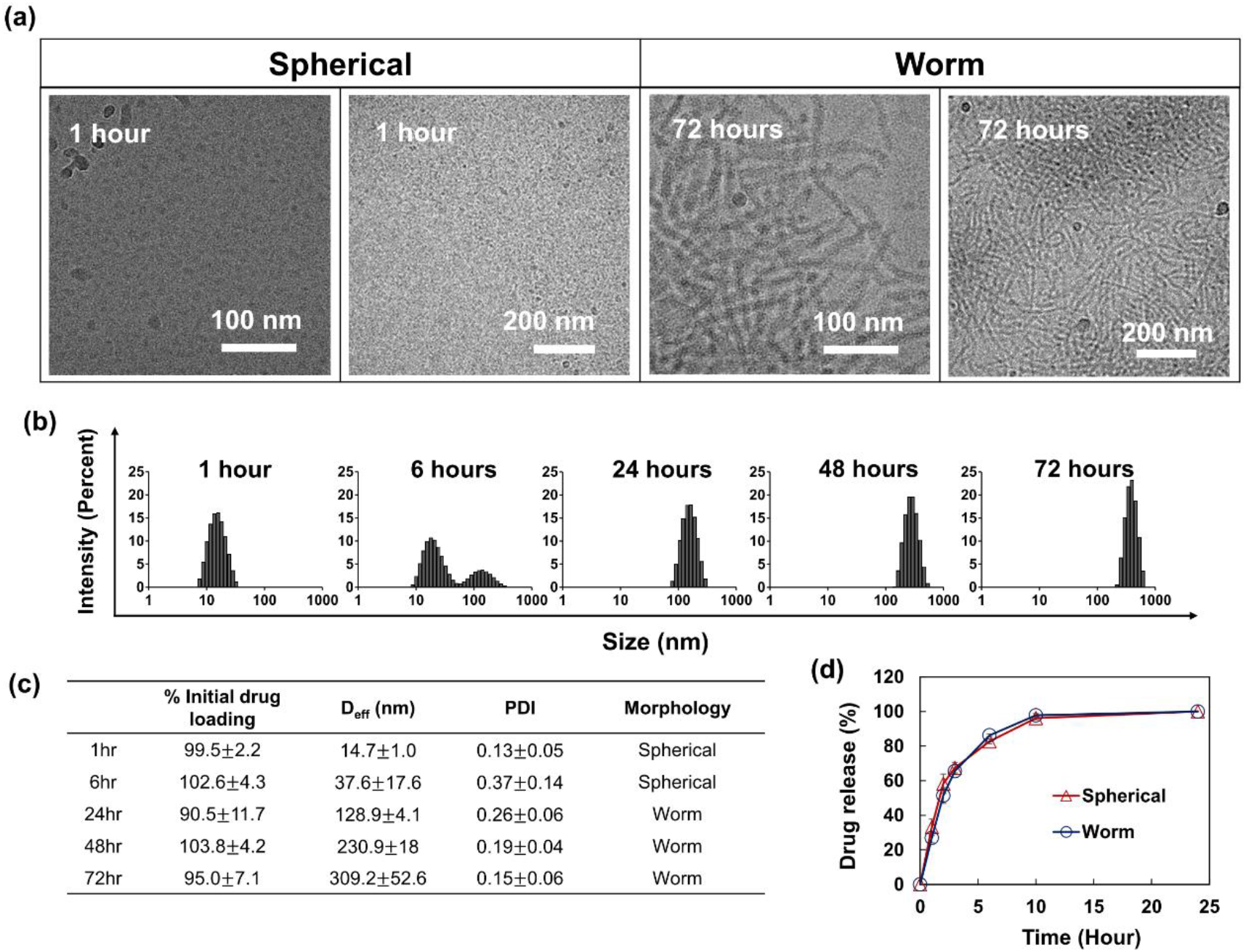
Preparation and characterization of olaparib-loaded micelles (polymer/drug ratio of 10/1) with different morphologies. (a) Cryo-TEM images of olaparib-loaded micelles at 1 hour and 72 hours after hydration. Change of (b) size distribution and (c) particle size, PDI, and drug concentration of olaparib-loaded micelles over time. (d) Olaparib release profile from sphere-and worm-like micelles. Olaparib-loaded micelles were prepared using the thin film hydration method. While keeping drug loading content, the olaparib micelles elongated until forming worm-like structure by 72 hours. Olaparib micelles did exhibit significant changes in the drug release profile. Errors shown correspond to three standard deviations of uncertainty.

### 2.8. Dependence of anti-tumor effect of a micellar drug on the micelle morphology

We evaluated the anti-tumor activity of olaparib against a tumor model (*BRCA1*-mutant human breast carcinoma xenograft MDA-MB-436; **Figure 9a**). Olaparib formulations in the spherical and worm-like micelles were administered intravenously (IV) at 10 mg/kg and 20 mg/kg, which correspond to ¼ and ½ of the drug maximum tolerated dose (MTD; 40 mg/kg) with a frequency of twice a week. Olaparib loaded in worm-like micelles showed significant anti-tumor efficacy by delaying tumor growth at the higher drug dose. These micelles also displayed a small effect at the lower dose, but differences were not statistically significant between drug treatment and saline (0.9 % NaCl) control groups. When the same drug was administered in spherical micelles, however, we observed significant antitumor activity at both drug doses (10 mg/kg and 20 mg/kg). Interestingly, the drug loaded in the spherical micelles was significantly more active than the drug loaded in the worm-like micelles, even at the same doses (**Figure 9a**).

**Figure 9.**
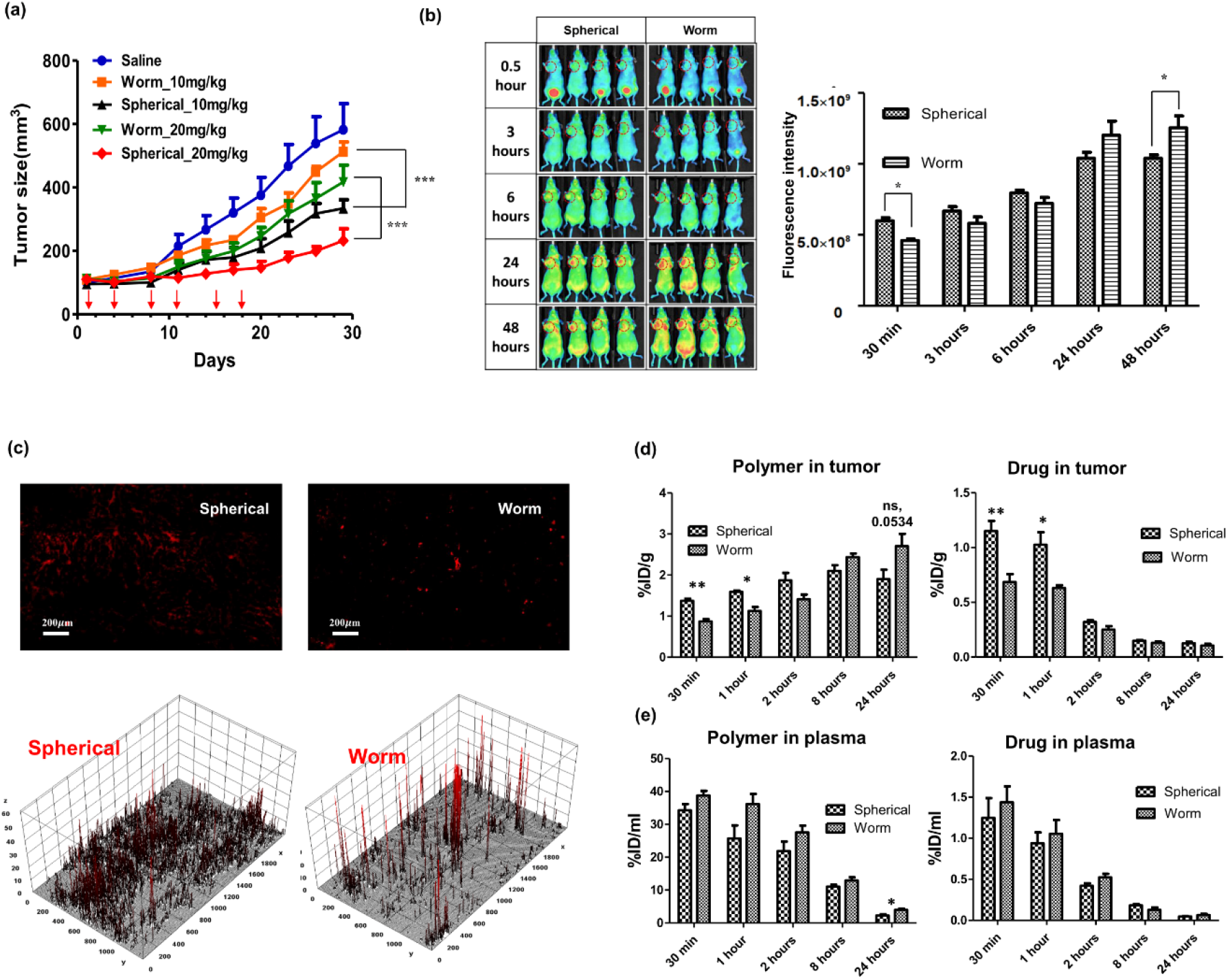
Antitumor activity, biodistribution, and PK of olaparib loaded micelles (polymer/drug ratio of 10/1) in a tumor model (BRCA1-mutant human breast carcinoma xenograft MDA-MB-436). **(a)** Tumor growth inhibition after 6 injections of olaparib-loaded micelles at ¼ and ½ of the drug MTD (40 mg/kg), delivered twice a week (shown by arrows). **(b)** Non-invasive fluorescent imaging of tumor model after a single injection of the AF-647 labeled micelles. **(c)** Visualization and 3D surface plots (analyzed in Image J) of the distribution AF-647 labeled micelles in tumor sections 1 hour after injection (see **Supplemental Figure S10** for blood vessel staining). **(d, e)** Quantification of the polymer and drug concentrations in **(d)** tumor and **(e)** plasma.

### 2.9. Effect of the micelle morphology on the PK and tumor distribution of the drug

AF-647 labeled micelles with different morphology were IV-injected into nude mice bearing MDA-MB-436 tumors; the distribution of the micelles was assessed by high-resolution fluorescent imaging of the tumor-bearing mice (**Figure 9b**). Micelles gradually accumulated at the tumor site over time for all formulations, although there were notable differences in accumulation kinetics (**Figure 9b**). In comparison to worm-like micelles, spherical micelles accumulated in the tumor more rapidly, at an early time (e.g., 30 min, 1 hour). However, after longer formation periods, the tumor accumulation of worm-like micelles gradually increased and exceeded the accumulation of spherical micelles after 24 hours. The AF-647–labeled micelles were visualized in tumor sections 1 hour after injection (**Figure 9c**). Both formulations accumulated in the tumor tissue but with different localization patterns. Overall, spherical micelles showed an even distribution and weak and scattered signal throughout the tumor tissue. Worm-like micelles, in contrast, were more localized and displayed a strong signal in a small area within tumor tissue (see Supplemental **Figure S10** for blood vessel staining).

Next, we carried out a PK study using tissue sampling with radiolabeled ^3^H-olaparib and AF-647–labeled micelles. We found differences in delivery to tumors of both the polymer and the drug when loaded in spherical or in worm-like micelles. At early time points (1 hour, 2 hours), we observed an increased accumulation of polymer administered as spherical micelles in the tumor compared to worm-like micelles (**Figure 9d**). This trend reversed over time, and accumulation of polymer in worm-like micelles increased steadily; however, these differences were not statistically significant. Similar trends were seen for the drug, with significantly higher tumor accumulation of olaparib in spherical micelles at 2 hours, compared to worm-like micelles. A maximum capacity (C_max_) of the drug was observed in the tumor at 0.5 hour (i.e., much earlier than the C_max_ of the polymer); this value was higher for spherical micelles compared to worm-like micelles (2.92 *μ*g/g vs. 1.75 *μ*g/g; p-value < 0.05) (**Table 1**). At later time points, this difference in drug accumulation for the two micellar morphologies became insignificant. Ultimately, the tumor AUC_all_ remained higher for the drug in spherical micelles due to the increased delivery of drug at an earlier stage (**Table 1**). The drug PK in the plasma were also similar for both spherical and worm-like micelles, whereas polymer PK in the plasma displayed a slight increase in C_max_ and statistically-significant difference in AUC for worm-like micelles compared to spherical micelles (**Figure 9e**). Therefore, both the polymer and the drug loaded in the spherical micelles distributed faster to the tumors, whereas distribution in the plasma during a period of 24 hours was comparable between the two morphologies, or even higher for worm-like micelles (see **Supplemental Table S3**).

**Table 1.** PK parameters of polymer and drug in tumor and plasma after a single injection of olaparib (10 mg/kg) in spherical (S) and worm-like (W) micelles (polymer/drug ratio of 10/1).

| Location | Parameter | Polymer |  |  | Drug |  |  |
| --- | --- | --- | --- | --- | --- | --- | --- |
|  |  | Spherical | Worm | Statistics | Spherical | Worm | Statistics |
| Tumor | T<sub>max</sub>(h) | 8.0 | 24.0 | - | 0.5 | 0.5 | - |
|  | C<sub>max</sub>(μg/mL) | 64.1 | 81.5 | ns | 2.9 | 1.7 | ** |
|  | AUC<sub>all</sub> (h × μg/mL) | 1466.4 | 1638.9 | ns | 13.8 | 11.4 | * |
| Plasma | T<sub>max</sub>(h) | 0.5 | 0.5 | - | 0.5 | 0.5 | - |
|  | C<sub>max</sub>(μg/mL) | 1105.5 | 1238.9 | ns | 3.2 | 3.7 | ns |
|  | AUC<sub>all</sub> (h × μg/mL) | 7999.6 | 9796.0 | * | 14.7 | 14.6 | ns |

<sup>a</sup> T<sub>max</sub> – time of the C<sub>max</sub>; C<sub>max</sub> – maximal observed concentration; AUC<sub>all</sub> – Area under the curve from the time of dosing to the time of the last observation.
<sup>b</sup> Significance S vs. W groups: \*p < 0.05, \*\*p < 0.01. See Supplemental Table S3 for detailed analysis of the polymer and drug exposure and clearance.

## 3. Discussion

From the perspective of nanomedicine manufacturing and clinical translation, it is critical to understand how nanoparticle morphology affects both polymer and drug biodistribution and therapeutic efficacy. Of particular relevance to this work, the distribution and clearance of polymeric micelles is reported to vary according to micellar morphology ^[1,12,13]^. However, most studies are limited to the study of *in vivo* fate of the nanoparticles themselves and do not include data on the fate of the encapsulated drug. Therefore, work elucidating the latter process and demystifying the correlation between micelle morphology, drug efficacy, and biodistribution is critical to enabling novel pharmacological discoveries.

Here, we investigated the elongation of poly(2-oxazoline)-based polymeric micelles from spherical to worm structure. Formation of the core-shell spherical micelles assembled into elongated worm-like large clusters was consistent across the cryo-TEM, DLS, and SANS observations. We found that the process of elongation is kinetically controlled and governed by a variety of conditions including the molecular mass of the polymer, polymer concentration, temperature, and ionic strength. The elongation process is also related to the drug loading of the micelles. As a general rule, when the amount of the added drug increased, the sphere-to-worm transition became slower or was completely inhibited. In view of this elongation and inhibiting process, we classified drugs as “worm-inhibiting” or “worm-permissive” groups.

The differences in the behavior of worm-inhibiting and worm-permissive drugs could be attributed to their specific interactions with the block copolymer, as characterized by the RD study. Considering that all the drugs used for the screening study are poorly soluble, we hypothesized that, in addition to hydrophobic interactions, other specific interactions between drug and polymer (e.g., hydrogen bonding or dipole-dipole interactions) may play a role in micelle stabilization. To assess the effects of these interactions on the micelle stability, we determined the changes of effective CMC* at various drug loadings; as micelles grew and elongated, the CMC* value decreased. This finding indicates increased stability of the worm-like morphology compared to the spherical micelles. The CMC* also gradually decreased as the drug loading amount increased. However, worm-inhibiting drugs decreased the CMC* of the spherical micelles upon loading to a greater extent and to the point where the transition to worm-like morphology became unfavored. The stabilization of the spherical micelle morphology implies that the worm-inhibiting drugs “prefer” spherical micelles as their environment. Based on this consideration, the use of the block copolymers that form spherical micelles offers the opportunity to increase loading capacity with respect to worm-inhibiting drugs. This seems to be the case of ABT-263, paclitaxel, and vismodegib, all of which display a two-fold increase in solubility in the T3 copolymer (forming the spherical micelles) relative to worm-forming T2 copolymer. No such difference was observed for worm-permissive drugs.

Overall, the knowledge and methodology acquired in this study enabled us to evaluate the role that morphology plays on the drug pharmacological performance of spherical and worm-like micelles. Morphology of polymeric micelles can be effectively controlled by tuning the type of polymer, molecular mass, and drug-loading ratio. Our findings suggest that morphology plays a critical role in the tumor accumulation behavior of micellular drug. Provided that spherical micelles are able to accumulate more rapidly, the tumor is exposed to micelles while they retain large amounts of drug. These results are consistent with other studies that show that worm-like micelles have longer circulation in the plasma ^[1, 13]^, which can lead to increased exposure and eventual accumulation in tumors. In contrast, worm-like micelles accumulate in tumors more slowly and only after they have released significant amounts of drug. As a result, less drug is delivered into the tumor by worm-like micelles. Although the overall systemic drug exposure is lower with spherical micelles, they result in more efficient tumor exposure and a two-fold increase in C_max_ partly due to the rapid tumor accumulation. Depending on the drug mechanism, both the exposure and/or C_max_ can drive pharmacological performance ^[14]^. Therefore, this result correlates with the increase in the anti-tumor effect of the drug administered by spherical micelles compared to worm-like micelles. The difference in anti-tumor activity is the result of the morphological transformations alone provided that all other parameters (e.g., drug regimen, doses, loading in the micelles, drug release rates, and the concentration and composition of the polymer) remained unchanged. The faster penetration of spherical micelles into tumors is likely due to their smaller size compared to worm-like micelles. These results confirm the decisive role that particle size plays in facilitating micelle entry and drug delivery to tumors ^[2, 12]^.

Other studies suggested the importance of nanoparticle shape in drug delivery; some of the benefits of leveraging particle nanoparticle morphologies include increased plasma circulation of rod-like nanoparticles ^[1, 13, 15]^, improved cell-binding-specificity of antibody-decorated to rod-like nanoparticles ^[16]^, and increased MTD of drugs in rod-like micelles ^[17]^. Our findings suggest that the dynamic character of the drug-micelle structure along with the micelle morphology play a pivotal role in pharmacological performance. In particular, our work indicates that drug release rates are important and that the changes in drug retention within the range of several hours could drastically influence pharmacological performance. This discovery sets our study apart from other nanoparticle studies, wherein drugs were either not included or were covalently conjugated.

Here, we demonstrated that spherical micelles facilitate systemic delivery of anticancer drugs to tumors when the drugs are loosely associated with the polymeric micelles. Our observations indicate that the polymeric micelle shape can be adjusted to a great extent by altering the block copolymer length, and that selection of a block length that favors the formation of spherical micelles can play a pivotal role in the formulation of pharmaceutical drugs. This is critical for improving drug loading, PK, and tumor distribution, and for ensuring long-term stability and reproducibility of formulations (e.g., particle size, polydispersity, and shape)—for which elongation over time is adverse.

## 4. Experimental Section

### Materials

All chemicals for polymer synthesis were purchased from Sigma-Aldrich (St. Louis, MO). Triblock copolymers of P[MeOx-*b*-BuOx-*b*-MeOx] with different degrees of polymerization of the blocks were synthesized by living cationic ring-opening polymerization, as described previously ^[18, 19]^. For the synthesis of polymers, methyl tri-fluoromethanesulfonate (MeOTf), 2-methyl-2-oxazoline (MeOx), and 2-n-butyl-2-oxazoline (BuOx) were dried by refluxing over calcium hydride (CaH_2_) under inert nitrogen gas and subsequently distilled. Blocks were sequentially synthesized using MeOTf as the initiator and terminated by piperidine. ^1^H NMR spectrum was acquired using an INOVA 400 system and analyzed using the MestReNova (11.0) software. Spectra were calibrated using MeOD solvent signals (4.78 ppm). Number-average molecular masses (M_n_) were determined using samples taken upon completion of each block and final copolymer, based on the ratio of ^1^H NMR signals of the initiator (CH_3_, 2.86-2.88, 2.97-3.03 ppm) and the repeating units of block backbone (C_2_H_4_, 3.40-3.70). Molecular mass distribution of the polymer was measured by Gel Permeation Chromatography (GPC) on a GPC-max VE2001 system; GPC data were used to determine the polymer polydispersity index (PDI).

Bortezomib, olaparib, etoposide, rapamycin, ABT-263, vismodegib, and paclitaxel were purchased from LC laboratories (Woburn, MA). BLZ945, RXDX-105, and PLX3397 were purchased from MedKoo Biosciences (Morrisville, NC). Resiquimod, AZD2014, and AZD8055 were purchased from Selleckchem (Houston, TX). All other chemicals were obtained from Fisher Scientific INC. (Fairlawn, NJ). All reagents used were of analytical grade.

MDA-MB-436 cells were obtained from UNC Lineberger Tissue Culture Facility and were cultured in a DMEM medium (Gibco 11965-092) supplemented with 10 % FBS and 1 % penicillin-streptomycin. The Cell Counting Kit (CCK-8) was purchased from Dojindo Molecular Technologies (Rockville, MD). Nude mice were purchased from UNC DLAM animal facility.

## Methods

### Preparation and characterization of POx-based polymeric micelles

POx micelles were prepared by the thin-film hydration method described previously ^[20]^. Briefly, stock solutions containing the polymer and the drugs in ethanol were mixed together at the pre-determined ratios and ethanol was then completely evaporated. The resulting thin film was hydrated with normal saline (0.9 % NaCl). The size distribution of micelle formulations resulting from various conditions was monitored over time using the DLS technique on a Zetasizer Nano ZS (Malvern Instruments Ltd., UK).

The amount of drug loaded in polymeric micelles was analyzed by reversed-phase high-pressure liquid chromatography (HPLC; Agilent Technologies 1200 series) performed using a Nucleosil C18, 5 *μ*m column (L × I.D. 250 mm × 4.6 mm). Each sample was diluted 50 times in a prescribed mobile phase (methanol/water volume percentage of 70 % / 30 %, with 0.01 trifluoroacetic acid for bortezomib; and acetonitrile/water volume percentage of 50 % / 50 %, with 0.01 trifluoroacetic acid for other drugs). Following this step, 10 *μ*L of diluted sample were injected into the HPLC while the flow rate was kept at 1.0 mL/min and the column temperature was kept at 40°C.

For TEM image acquisition, we used a LEO EM910 TEM operating at 80 kV (Carl Zeiss SMT Inc., Peabody, MA). Diluted samples were deposited onto a copper grid containing a carbon film and stained with 1% uranyl acetate before imaging. For Cryo-TEM image acquisition, cryo-grids were prepared by rapid immersion into an ethane/propane mixture (40 % / 60 %) using a Vitrobot Mark IV (Thermo Fisher Scientific) set to 22 °C and 95 % humidity. C-Flat TEM grids (ProtoChips) were rendered hydrophilic by indirect oxygen plasma cleaning (25 % oxygen, 75 % argon) using a Tergeo-EM system (PIE Scientific LLC). Cryo-grids were imaged using a 200 kV Talos Artica G3 (Thermo Fisher Scientific) equipped with a Gatan K3 direct-electron detector (Gatan).

### Screening of micelle elongation

The effect of various conditions including polymer concentration (1.25∼80 mg/mL), temperature (4 °C, 25 °C, and 37 °C), and salt concentration (up to ∼4 % NaCl) on micelle elongations were investigated by monitoring the change of particle size and size distribution over time using DLS (Zetasizer Nano ZS; Malvern Instruments Ltd., UK). To examine drug-loading effects on micelle elongation, drugs were formulated into polymeric micelles at predetermined drug loading ratios (polymer/drug w/w ratios of 10/0, 10/1, 10/2, 10/4 and 10/8); the change of particle size and size distribution were monitored over time. The morphology of each formulation was classified into three categories based on particle size and size distribution: (1) spherical micelles (S), (2) mixtures of spherical and worm micelles (W+S), and (3) worm micelles (W). If micelles were destabilized and precipitated, samples were classified as precipitate (P).

### Small Angle Neutron Scattering (SANS)

To perform SANS measurements, blank and drug-loaded micelle solutions were prepared in D_2_O containing 10 mg/mL of T2. All samples were degassed before SANS measurements; these were examined using the NG7 and NGB30 systems ^[21]^ at the National Institute of Standards and Technology Center for Neutron Research (NCNR). Data collection was performed at 25°C. A neutron wavelength *λ* of 6 Å and three sample-to-detector distances were used. A high resolution lens configuration (λ = 8.09 Å) provided the lower scattering angle data. SANS data were collected in the range of momentum transfer 0.002 < *q* (Å ^-1^) < 0.55, where *q* is defined as follows:

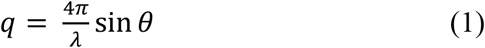

where the scattering angle is given by 2*θ*. Data were reduced and analyzed using the SANS macro routines and analysis packages, respectively, developed for IGOR at the NCNR ^[22]^. Raw counts were normalized to a common monitor count and corrected for empty cell counts, ambient background counts, and nonuniform detector response. Scattered intensities were corrected to remove incoherent scattering (e.g., from the buffer and from hydrogen) using a flat background subtraction based on the lowest intensity recorded at high *q* (for each sample).

SANS data were measured for solutions containing a constant concentration of polymer and increasing amounts of drug (polymer/drug w/w ratios of 10/0, 10/1, 10/2, and 10/4; **Figure 4**). An upturn in scattered intensities for the blank and lower drug concentration micelle solutions was observed at *q* lower than ≈ 0.015 Å^-1^. This is characteristic of larger clusters or aggregates and is consistent with cryo-TEM and light scattering observations. The average size of these larger objects is beyond the angular range measured by SANS. Data were fit with the following functional form:

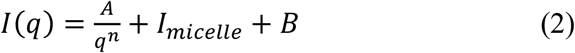

where the first term describes Porod scattering from clusters of micelles, B is the incoherent background, and I_micelle_ denotes the scattering from individual T2 spherical micelles (blank or drug loaded). With increasing drug loading ratio, single micelles are stabilized, and a flattening of the lower-q upturn is observed. Contributions to scattering from the individual micelles were modelled as follows:

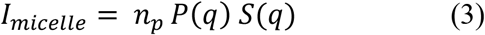

where *n*_*p*_ is number density of micelles with an effective sphere diameter *σ. P(q)* was modelled using a polydisperse sphere where the ratio between the core radius and the total micellar radius is held constant ^[23]^. *P(q)* is the particle form factor averaged over the particle size distribution:

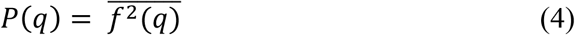

The single particle scattering amplitude is an average over a Schulz distribution of radii (see equations 32-37 in reference ^[23]^), where:

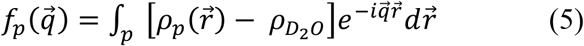

*ρ* is the particle SLD and *r* is the distance between particles. An average structure factor approximation was used:

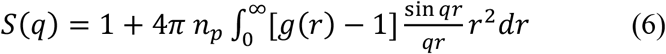

where g(r) is the pair correlation function defined by a hard sphere interparticle potential. The Ornstein-Zernicke equation was solved using a Percus-Yevick closure as described previously ^[22,24]^.

### Light scattering intensity

The effective CMC* of poly(2-oxazoline)-based polymeric micelles was determined by DLS (wavelength 663nm). The light scattered by the samples was detected by an avalanche photodiode at a backscattering angle of 173 degrees. The intensity of scattered light (derived count rates, kilo-counts per second (kcps)) was obtained for blank and drug loaded micelles (1 mg/mL), diluted in distilled water to varying concentrations. Above a given polymer concentration, the intensity of light scattering proportionally decreased as polymer concentration decreased; above this concentration threshold, there were no significant changes in the intensity of light scattering. This point of deflection (e.g., slope change) in the intensity of light scattering was defined as effective CMC* ^[25]^.

### Reichardt’s dye (RD)

RD and drug-loaded polymeric micelles were prepared using a thin-film hydration method. The polymer, drugs, and Reichardt’s dye were mixed in ethanol at pre-determined ratios (10/0.5/0.5, 10/1/0.5, 10/2/0.5 and 10/4/0.5), after which ethanol was completely evaporated from the mixtures. The resulting thin film was subsequently hydrated using normal saline (0.9 % NaCl). After 1 hour stabilization at RT, the UV-vis spectra of each sample were recorded. The maximum absorption wavelength and the absorbance intensity values of RD were analyzed.

### Molecular dynamics simulation (MD)

MD simulations were performed in GROMACS 2020 using an OPLS-AA force field. Polymer and drug were packed randomly in a cubic box and surrounded with water and salt. The structure was annealed to 333 K for the first 50 ns of the simulation to equilibrate the system. The system was then cooled back to 300 K for at least 100 ns. Partial density was computed using the “density” module in GROMACS.

### *In vitro* drug release

Drug release profiles were obtained using the membrane dialysis method described elsewhere ^[19, 26]^. First, the olaparib loaded micelles were diluted in phosphate-buffered saline (PBS) to achieve a drug concentration of 0.1 mg/mL. Then, the diluted micelle solutions were transferred to floatable Slide-A-Lyzer MINI dialysis devices (100 *μ*L capacity, 3.5 kDa MWCO) and suspended in 30 mL of PBS supplemented with 10 % fetal bovine serum (FBS) (to ensure sink conditions). Four devices were used for every time point. At predetermined time points, the samples were collected from the devices and the remaining drug amounts were quantified by HPLC. Drug release profiles were constructed by plotting the percentage released over time.

### Estimation of Maximum Tolerated Dose (MTD)

A dose-escalation study was used to estimate the MTD before *in vivo* efficacy studies. Healthy nude mice (8 weeks old, female) were divided into groups of four, with each group subjected to increasing doses of drugs. Olaparib PM (20, 40, and 60 mg/kg) and normal saline (0.9 % NaCl) were injected intravenously using a q4d x 6 regimen. Mice were examined every other day to monitor signs of toxicity, including hunched posture, rough coat, and body weight changes (loss of 15 % or greater) in accordance to approved Institutional Animal Care and Use Committee (IACUC) protocols.

### Tumor inhibition

Female athymic nude mice (6–8 weeks) were subcutaneously inoculated in the right flank with MDA-MB-436 cells (5 × 10^6^ cells in 100 uL of 1:1 mix of PBS and BD Matrigel). When the tumor sizes reached ca. 100 mm^3^, mice were randomized into 5 groups, and the animals (n=5) received the following injections by group: (1) Normal saline (i.v.; q4d x 6); (2) 10 mg/kg of spherical-shaped olaparib PM (i.v.; q4d x 6); (3) 10 mg/kg of worm-shaped olaparib PM (i.v.; q4d x 6); (4) 20 mg/kg of spherical0shaped olaparib PM (i.v.; q4d x 6); or (5) 20 mg/kg of worm-shaped olaparib PM (i.v.; q4d x 6). Tumor volume was calculated using the formula (*length x width*^2^)/2. Tumor growth and body weight of the mice were monitored for 1 month.

### *In vivo* particle distribution

Nude mice bearing MDA-MB-436 tumors were IV-injected with a single dose of spherical or worm-shaped olaparib PM micelles (10 mg/kg) containing a AF-647 conjugated polymer (2.5 % of total polymer). After 1 hour from sample injections, mice were euthanized. Tumors were then collected and frozen in a cryo-mold filled with an optimal cutting temperature (OCT) compound (Tissue-Tek; Sakura Finetek, Leiden, The Netherlands). Tumor cryosections were obtained at a thickness of 10 *μ*m and tumor vasculatures were stained with anti-CD31 (Abcam, Cambridge, UK) with assistance from Translational Pathology Laboratories. Stained slides were digitally acquired using an Aperio ScanScope XT (Aperio). Polymer and CD31 intensity at tumor sections were visualized by 3D surface plot analysis (ImageJ). The X and Y axes represent the area of the tumor sections, whereas the Z axis represents the polymer (Red) and CD31 (Green) intensities.

### Pharmacokinetic (PK) and biodistribution studies

Female athymic nude mice (6-8 weeks) were subcutaneously inoculated in the right flank with MDA-MB-436 cells (5 × 10^6^ cells in 100 *μ*L of 1:1 mix of PBS and BD Matrigel). PK studies proceeded once tumor sizes reached ca. 200 mm^3^.

For drug PK studies, mice were administered a single dose of spherical or worm-shaped olaparib-loaded micelles (10 mg/kg) containing ^3^H-labeled olaparib (200 *μ* Ci/kg) *via* tail vein. At predetermined time points (0.5, 1, 2, 8, 24, and 48) hours, each group of mice (n=3∼4) was euthanized; samples (plasma and tissues) were collected to analyze the olaparib concentration by measuring the radioactivity. A portion of plasma was diluted in mixture of Soluene-350 and isopropyl alcohol (1:1 ratio) and incubated at 60 °C for 2 hours. Tissues were weighed and homogenized using a homogenizer Tissuemiser (Fisher Scientific, Pittsburgh, PA) and Soluene-350. A portion of the blood or tissue homogenate was placed in a scintillation vial and mixed with 10 mL of scintillation cocktail (Ultima Gold). Radioactivity was measured using a liquid scintillation counter, Tricarb 4000 (Packard, Meriden, CT).

For polymer PK studies, mice were administered a single dose of spherical or worm-shaped olaparib-loaded micelles (10 mg/kg) containing an AF-647–conjugated polymer (2.5 % of total polymer) *via* tail vein. Plasma was collected and diluted in saline (0.9 % NaCl) and analyzed using fluorescence microscopy. Organs were collected and homogenized in a mixture of 2 mg/mL of collagenase and Ripa buffer (1:1 ratio). Polymers in homogenate were extracted using MeOH, and their fluorescence intensity was analyzed using a SpectraMax M5 plate reader (Molecular Devices, Sunnyvale, CA) (excitation = 633 nm, emission = 670 nm).

PK data were analyzed using Phoenix Winnonlin modeling software. Noncompartmental analysis (NCA) was performed using sparse sampling; NCA parameters were used as initial estimates for model development. AUC and residual plot analyses were utilized as criteria for determining the best model for the polymers and the drug.

### Statistical analysis

Statistical comparison of data for tumor inhibition was done using a two-way analysis of variance (ANOVA) and followed by Bonferroni post-tests for multiple comparison. Comparison of particle accumulation in tumor and PK were done by unpaired t-test. Statistical analyses were performed using the GraphPad Prism software (***p< 0.001).

## Acknowledgements

This work was partially supported by the National Cancer Institute (NCI) Alliance for Nanotechnology in Cancer (U54CA198999, Carolina Center of Cancer Nanotechnology Excellence). Animal Studies and IVIS imaging were performed within the UNC Lineberger Animal Studies Core (ASC) Facility at the University of North Carolina at Chapel Hill. The UNC Lineberger Animal Studies Core is supported, in part, by an NCI Center Core Support Grant (CA16086) to the UNC Lineberger Comprehensive Cancer Center. Access to the NGB 30m SANS instrument was provided by the Center for High Resolution Neutron Scattering, a partnership between the National Institute of Standards and Technology and the National Science Foundation under Agreement No. DMR-2010792. Cryogrids were prepared and imaged at the UNC Chapel Hill CryoEM Core. We would like to thank the University of North Carolina at Chapel Hill and the Research Computing group for providing computational resources and support that have contributed to these research results. We thank C. Santos, M. Ross, and A. Valdivia at MP1U of UNC for helping with the intravenous/intraperitoneal injections. TEM was performed by A. Shankar Kumbhar at the Chapel Hill Analytical and Nanofabrication Laboratory (CHANL), a member of the North Carolina Research Triangle Nanotechnology Network (RTNN), which is supported by the NSF (grant ECCS-1542015) as part of the National Nanotechnology Coordinated Infrastructure (NNCI). We acknowledge the support of the National Institute of Standards and Technology, U.S. Department of Commerce, in providing the neutron research facilities used in this work. SCMT is very grateful for the general support and useful discussions with the staff at the NIST Centre for Neutron Research (NCNR), and funding from CNS NIST Cooperative Agreement 70NANB12H239.

## Disclaimer

Certain commercial equipment, instruments, suppliers are identified to foster understanding. This does not imply recommendation or endorsement by the National Institute of Standards and Technology, nor does it imply that the materials or equipment identified are necessarily the best available for the purpose.

