## Supplementary Materials for "Drug-dependent morphological transitions in spherical and worm-like polymeric micelles define stability and pharmacological performance of micellar drugs"

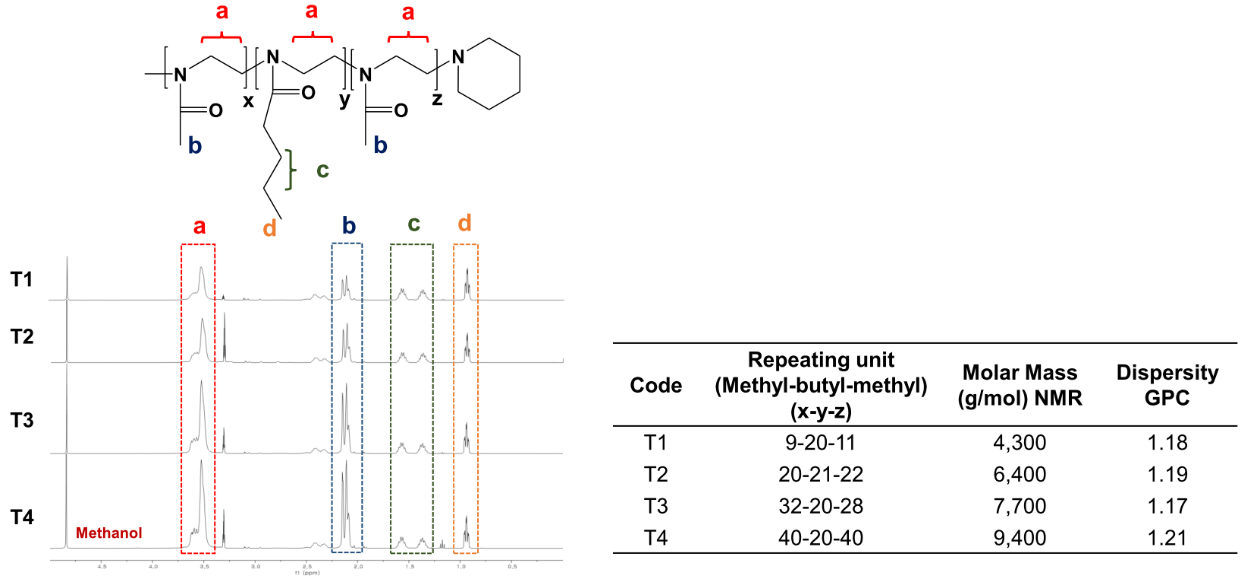

**Figure S1.** Structure, NMR spectra, and molecular characteristics of the poly(2-oxazoline)-based triblock copolymers used in this study. (Dispersity = polydispersity index)

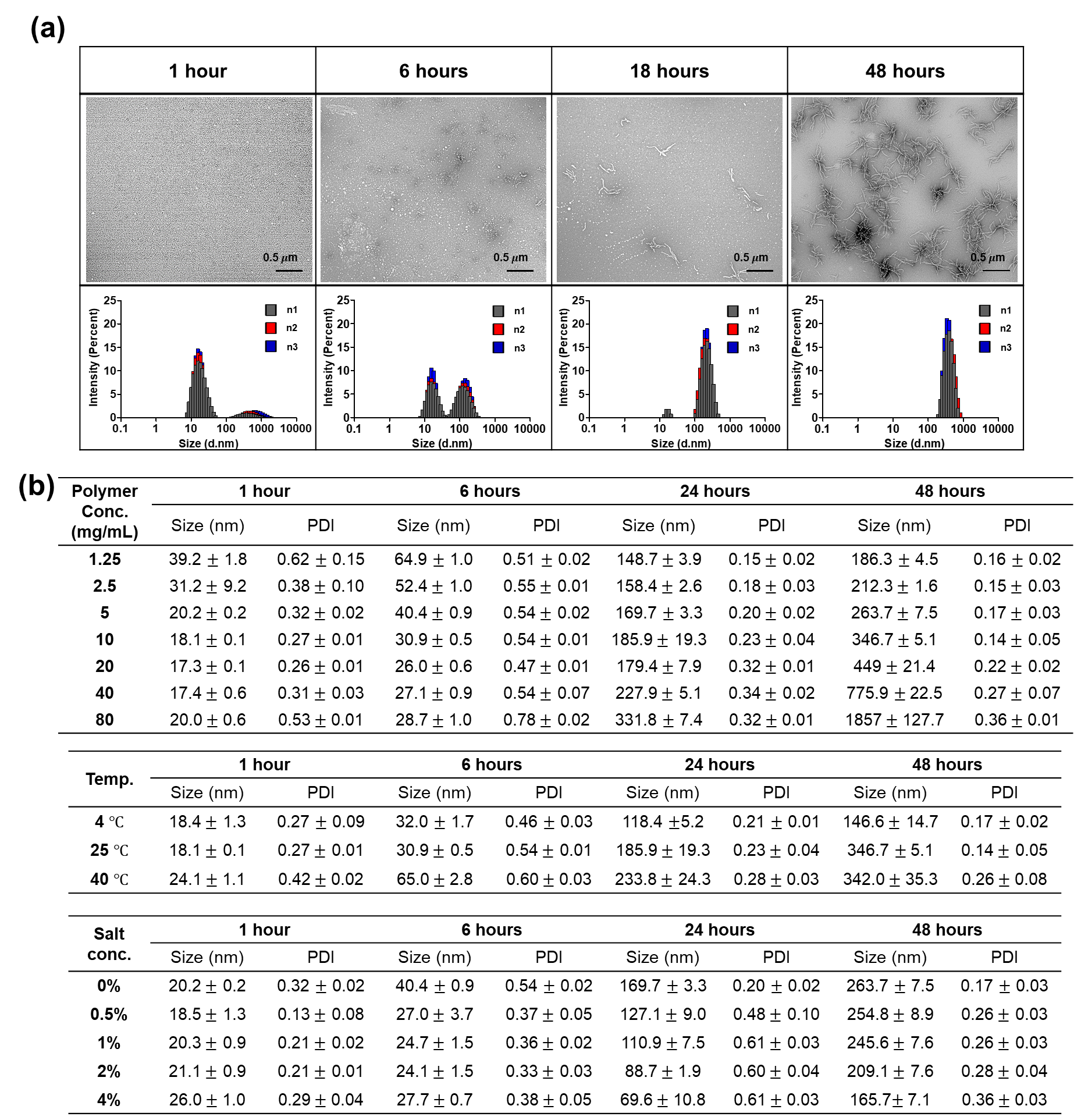

**Figure S2. (a)** Evolution of the particle morphology and size distribution of the T2 polymeric micelles in distilled water (10 mg/mL, 25^o^C), visualized by TEM. **(b)** Particle size and PDI of the T2 polymeric micelles under various conditions, to examine: the effect of polymer concentration (in distilled water and at 25℃), temperature (10 mg/mL polymer concentration in distilled water) and ionic strength (5 mg/mL polymer concentration at 25℃). Elongation was favored by increasing the concentration and temperature and inhibited by increasing the ionic strength. The errors shown correspond to three standard deviation of uncertainty.

Our findings suggest that (1) the elongation process is kinetically controlled, taking many hours to achieve the worm-like state, and that (2) the resulting morphologies depend on the hydrophilic block lengths (see **Figure S2a** and also main **Figure 1**). On one hand, with the T1 triblock copolymer where the length of the hydrophilic block was short (10 r.u. (repeating unit)), micelles grew 3-dimensionaly but did so anisotropically. On the other hand, when the length of the hydrophilic block was longer at 20 r.u. (as seen in the T2 particles), micelles elongated in a 1-D manner resulting in longer, worm-like structures. In the case of the T3 and T4 polymers, with even longer hydrophilic blocks (30 r.u. and 40 r.u., respectively), some of the micelles elongated in a 1-D manner but many structures remained spherical in both formulations even after 72 hours. The T1 and T2 copolymers, with the shortest hydrophilic blocks, likely cannot ensure sufficient repulsion in the shell to counteract transition to worm-like morphology, which is accompanied by the increase of the density of the chains in the shell. As the length of the hydrophilic chain increased (e.g., from T2 to T3 and T4), the repulsion of the hydrophilic chains also increased and favored the spherical micelles, where the surface area per one hydrophilic chain is greater. While the worm-like morphology is more favorable in a shorter hydrophilic block, it is partly inhibited in longer hydrophilic blocks.

We also investigated micelle elongation with respect to the polymer concentration, temperature, and salt concentration (**Figure S2b**). These parameters were used to effectively tune the elongation process. We observed that, when the polymer concentration increased, the elongation rate also increased potentially due to the increased rate of micelle collision resulting in polymer-polymer interactions and elongation. When the temperature increased, the elongation rate increased possibly due to the increase in hydrophobic interactions. Generally, the contact between the hydrophobic core and water should promote micelle growth. At increased temperature, however, the exposure of the hydrophobic core to water molecules becomes less favorable, resulting in shrinkage of the core surface area. This could yield favorable conditions for micelle elongation—both kinetically and thermodynamically. As the salt concentration increased, the micelle elongation rate decreased. The addition of salt likely dehydrates the polymer chains in the hydrophobic core, increasing the core density, decreasing the surface area and favoring the formation of spherical micelles. Altogether, these parameters could provide control over micelle stability and morphology, yielding a more predictable behavior during the formation process.

**Table S1.** Loading parameters (LC, LE)^a^, DLS (Dynamic light scattering) particle size (D_eff_), and polydispersity (PDI) of the micelles at different drug loadings (polymer/drug w/w ratios of 10/1, 10/2, 10/4 and 10/8). The errors shown represent three standard deviation of uncertainty.

| **Drug** | | **LC**_max_ **(%)** | **LE**_max_ **(%)** | **10/1** | | **10/2** | | **10/4** | | **10/8** | |
| --- | --- | --- | --- | --- | --- | --- | --- | --- | --- | --- | --- |
|  |  |  |  | **D_eff_ (nm)** | **PDI** | **D_eff_ (nm)** | **PDI** | **D_eff_ (nm)** | **PDI** | **D_eff_ (nm)** | **PDI** |
| T2 | Bortezomib | 43.4 ± 1.3 | 97.5 ± 3.0 | 16.2 ± 1.2 | 0.16 ± 0.06 | 15.8 ± 0.2 | 0.15 ± 0.02 | 16.3 ± 0.3 | 0.08 ± 0.02 | 22.6 ± 1.8 | 0.10 ± 0.06 |
|  | Olaparib | 27.3 ± 1.0 | 93.9 ± 4.8 | 14.7 ± 1.0 | 0.13 ± 0.05 | 18.3 ± 1.5 | 0.19 ± 0.06 | 20.5 ± 3.3 | 0.13 ± 0.01 | N/A | |
|  | Etoposide | 42.0 ± 2.9 | 91.0 ± 10.6 | 15.5 ± 0.5 | 0.08 ± 0.02 | 18.2 ± 2.0 | 0.14 ± 0.02 | 18.2 ± 1.9 | 0.17 ± 0.04 | 25.3 ± 4.6 | 0.09 ± 0.02 |
|  | Resiquimod | 43.4 ± 1.0 | 95.7 ± 3.8 | 21.4 ± 4.3 | 0.19 ± 0.02 | 20.5 ± 3.9 | 0.19 ± 0.01 | 17.6 ± 0.8 | 0.19 ± 0.04 | 19.9 ± 1.2 | 0.25 ± 0.03 |
|  | BLZ945 | 42.5 ± 1.9 | 92.7 ± 7.5 | 23.7 ± 2.0 | 0.28 ± 0.03 | 20.7 ± 1.9 | 0.15 ± 0.05 | 20.3 ± 0.9 | 0.12 ± 0.03 | 27.5 ± 2.6 | 0.03 ± 0.01 |
|  | Rapamycin | 26.5 ± 0.8 | 90.1 ± 3.8 | 22.8 ± 2.6 | 0.22 ± 0.06 | 19.8 ± 1.0 | 0.19 ± 0.03 | 22.0 ± 1.4 | 0.04 ± 0.03 | N/A | |
|  | RXDX-105 | 27.6 ± 1.9 | 95.5 ± 8.8 | 24.3 ± 2.5 | 0.15 ± 0.04 | 20.3 ± 1.8 | 0.19 ± 0.11 | 24.1 ± 3.2 | 0.19 ± 0.12 | N/A | |
|  | AZD2014 | 28.4 ± 1.1 | 99.4 ± 5.5 | 31.1 ± 2.7 | 0.15 ± 0.08 | 20.1 ± 0.5 | 0.25 ± 0.03 | 19.2 ± 0.3 | 0.11 ± 0.04 | N/A | |
|  | AZD8055 | 44.3 ± 1.0 | 99.3 ± 4.0 | 20.4 ± 3.0 | 0.25 ± 0.08 | 19.1 ± 0.2 | 0.16 ± 0.01 | 21.8 ± 0.6 | 0.11 ± 0.03 | 28.8 ± 0.5 | 0.03 ± 0.02 |
|  | PLX3397 | 27.8 ± 1.5 | 96.7 ± 7.3 | 18.6 ± 0.5 | 0.21 ± 0.03 | 22.5 ± 1.5 | 0.25 ± 0.08 | 54.9 ± 0.8 | 0.17 ± 0.03 | N/A | |
|  | Vismodegib | 27.3 ± 1.7 | 95.6 ± 5.9 | 18.1 ± 0.8 | 0.06 ± 0.01 | 19.1 ± 1.3 | 0.05 ± 0.02 | 24.8 ± 2.1 | 0.08 ± 0.01 | N/A | |
|  | Paclitaxel | 27.1 ± 2.4 | 92.9 ± 11.2 | 16.2 ± 0.6 | 0.03 ± 0.01 | 18.6 ± 0.9 | 0.07 ± 0.01 | 18.0 ± 0.5 | 0.12 ± 0.01 | N/A | |
| T3 | Bortezomib | 43.5 ± 1.2 | 97.9 ± 2.7 | 18.5 ± 2.2 | 0.15 ± 0.01 | 17.7 ± 2.8 | 0.12 ± 0.01 | 19.9 ± 2.9 | 0.12 ± 0.01 | 107.6 ±8.3 | 0.06 ± 0.04 |
|  | Olaparib | 28.1 ± 0.9 | 97.5 ± 4.5 | 18.0 ± 3.0 | 0.11 ± 0.05 | 17.4 ± 0.6 | 0.14 ± 0.02 | 20.2 ± 2.7 | 0.16 ± 0.03 | N/A | |
|  | Etoposide | 43.2 ± 0.5 | 95.0 ± 1.9 | 17.5 ± 1.2 | 0.09 ± 0.02 | 18.2 ± 2.1 | 0.10 ± 0.04 | 20.3 ± 2.1 | 0.08 ± 0.02 | 33.1 ± 1.6 | 0.13 ± 0.01 |
|  | Resiquimod | 42.3 ± 2.9 | 91.9 ± 11.2 | 22.1 ± 6.3 | 0.10 ± 0.03 | 19.9 ± 3.3 | 0.20 ± 0.01 | 21.3 ± 2.8 | 0.23 ± 0.01 | 26.4 ± 2.7 | 0.21 ± 0.03 |
|  | BLZ945 | 42.8 ± 1.9 | 93.7 ± 7.2 | 19.3 ± 0.4 | 0.09 ± 0.04 | 23.7 ± 3.0 | 0.13 ± 0.07 | 40.2 ± 6.5 | 0.16 ± 0.03 | 87.9 ± 1.4 | 0.09 ± 0.03 |
|  | Rapamycin | 42.6 ± 2.2 | 92.8 ± 8.1 | 19.9 ± 1.0 | 0.10 ± 0.02 | 20.8 ± 0.7 | 0.07 ± 0.05 | 29.2 ± 1.5 | 0.08 ± 0.05 | 53.5 ± 3.4 | 0.19 ± 0.06 |
|  | RXDX-105 | 42.5 ± 2.1 | 92.8 ± 7.9 | 33.5 ± 1.0 | 0.18 ± 0.02 | 36.6 ± 0.5 | 0.17 ± 0.05 | 32.8 ± 0.4 | 0.13 ± 0.02 | 81.4 ± 7.2 | 0.22 ± 0.02 |
|  | AZD2014 | 27.8 ± 1.3 | 96.3 ± 6.3 | 19.2 ± 0.5 | 0.13 ± 0.04 | 19.5 ± 0.3 | 0.13 ± 0.02 | 22.3 ± 1.9 | 0.13 ± 0.06 | N/A | |
|  | AZD8055 | 43.3 ± 2.1 | 95.7 ± 7.9 | 24.4 ± 3.2 | 0.22 ± 0.03 | 23.1 ± 0.7 | 0.17 ± 0.06 | 34.6 ± 0.4 | 0.25 ± 0.02 | 77.2 ± 15.9 | 0.23 ± 0.02 |
|  | PLX3397 | 43.8 ± 1.5 | 97.4 ± 6.2 | 26.9 ± 0.3 | 0.20 ± 0.01 | 23.6 ± 0.8 | 0.11 ± 0.02 | 31.1 ± 1.3 | 0.14 ± 0.04 | 88.7 ± 1.1 | 0.19 ± 0.03 |
|  | Vismodegib | 42.8 ± 1.7 | 96.3 ± 3.9 | 20.6 ± 0.5 | 0.11 ± 0.01 | 21.1 ± 2.5 | 0.08 ± 0.04 | 27.1 ± 3.9 | 0.04 ± 0.05 | 41.4 ± 4.3 | 0.11 ± 0.03 |
|  | Paclitaxel | 43.0 ± 1.3 | 94.4 ± 4.9 | 20.8 ± 1.4 | 0.11 ± 0.02 | 24.0 ± 0.3 | 0.09 ± 0.04 | 33.6 ± 0.9 | 0.20 ± 0.02 | 54.1 ± 5.9 | 0.15 ± 0.05 |

^a^ Loading capacity, $LC \%=\frac{mass drug loaded}{mass drug loaded+mass polymer used}\times100$; Loading efficiency, $LE \%=\frac{mass drug loaded}{mass drug added}\times100$

PLX3397 even at low loadings formed unstable micelle formulations, which precipitated without apparent formation of worms after 24 hours. In other cases, precipitation was observed only after high drug loadings.

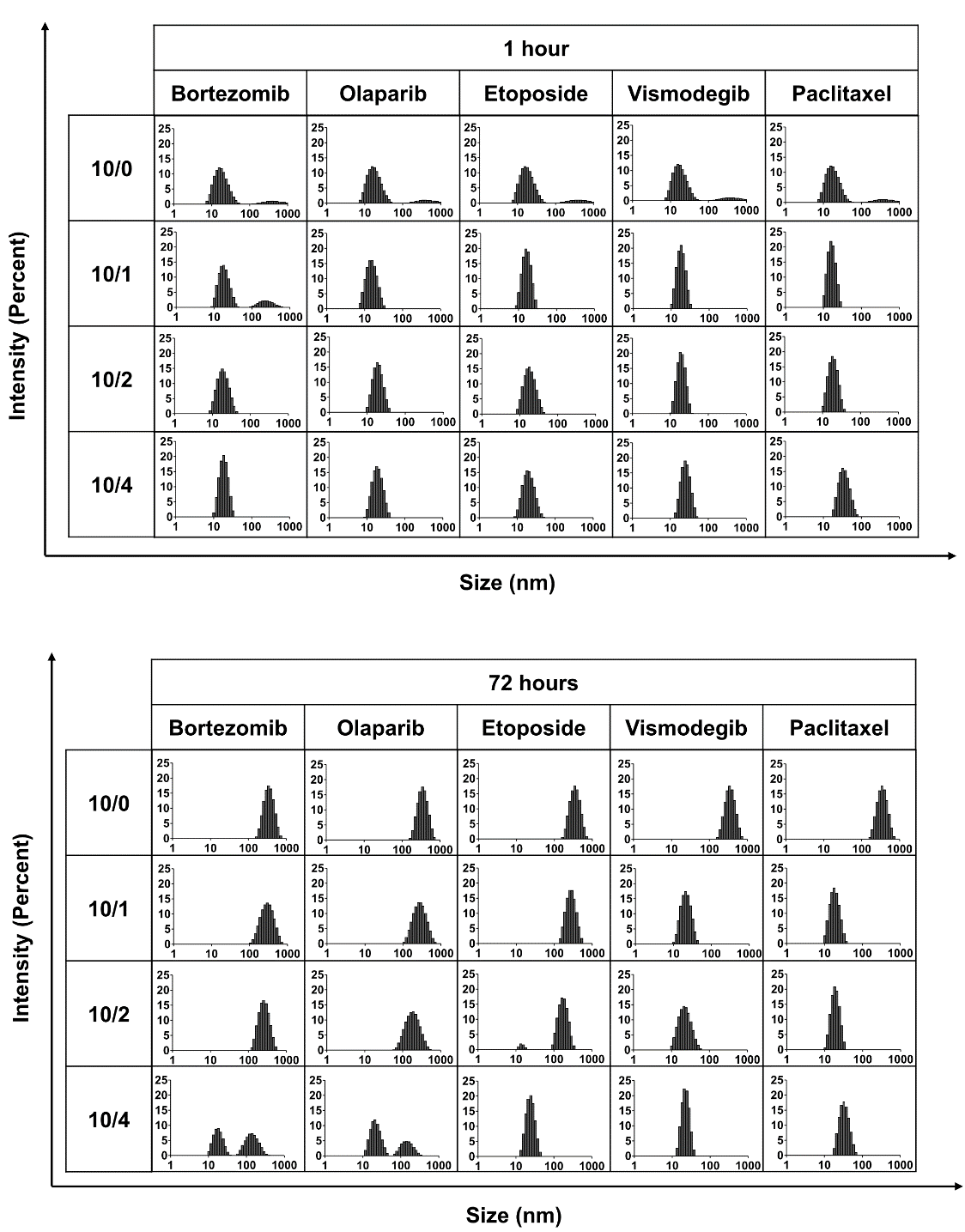

**Figure S3.** Change in size distribution of T2-based polymeric micelles at various drug loading ratios over time (polymer/drug ratios of 10/1, 10/2, and 10/4). Representative data for aggregation of T2. Small spherical micelles, with the particle size of ca. 20–30 nm, are observed at short incubation times, both in the drug-free micelle solutions and/or in the presence of worm-inhibiting drugs at high drug loadings. The observed increase over time in particle size, up to several hundred nm, is indicative of worm formation.

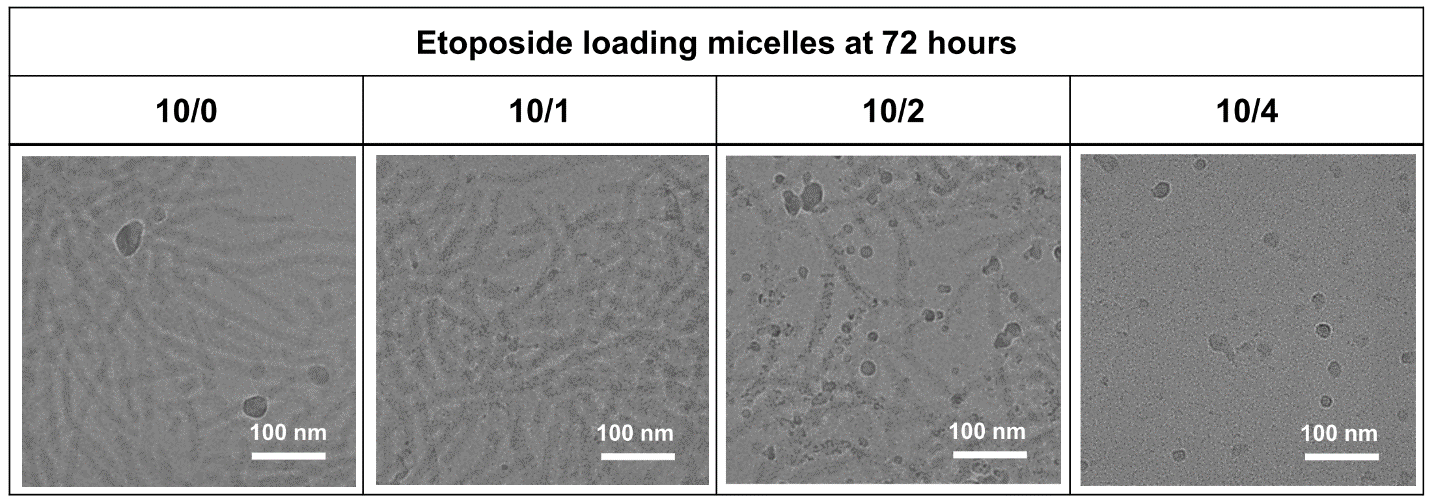

**Figure S4.** Cryo-TEM images of T2 aggregates in the presence and absence of various amounts of etoposide at room temperature, after 72 hours. T2 concentration 10 mg/mL in saline (0.9 % NaCl).

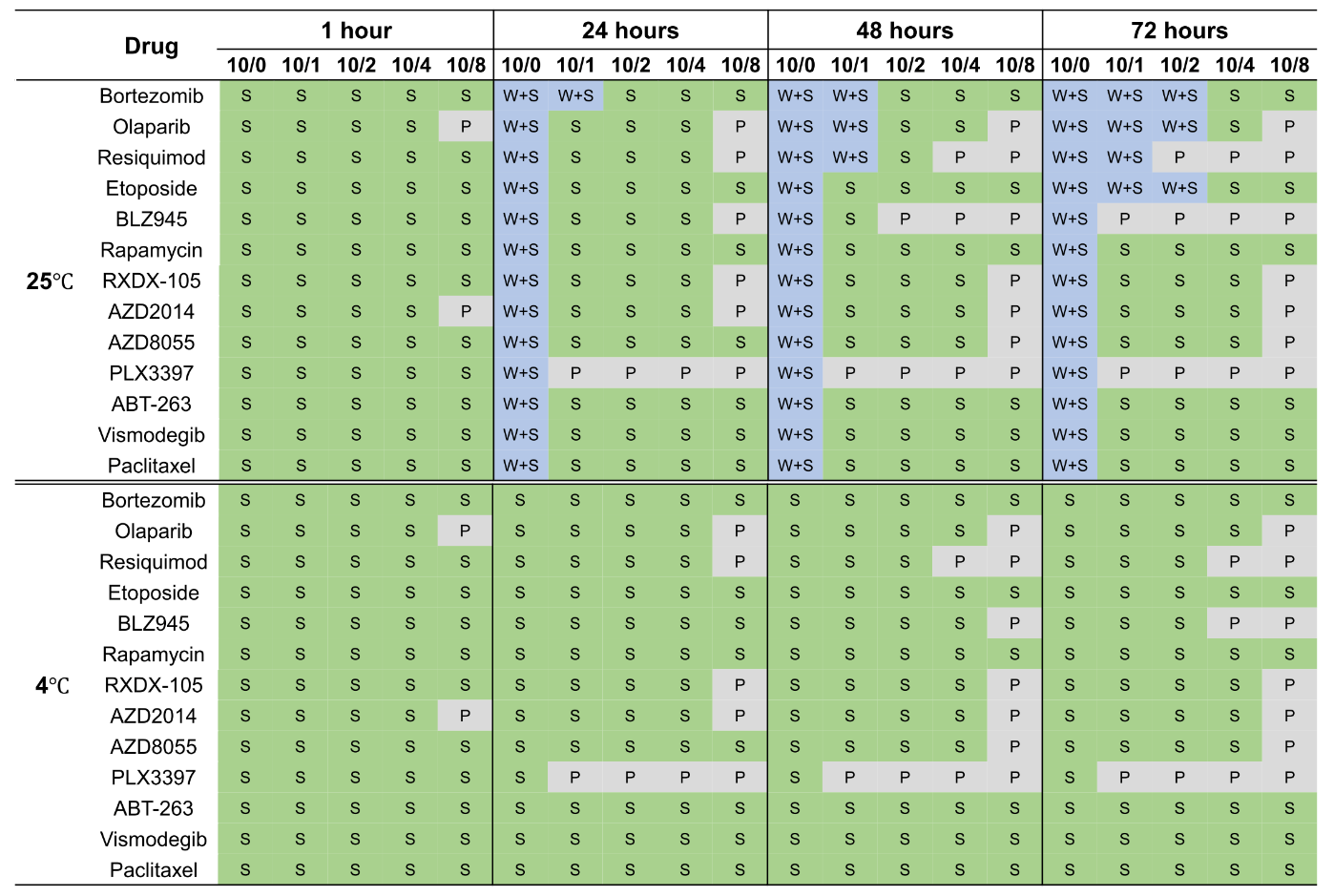

**Figure S5.** Monitoring the sphere-to-worm transition process of T3-based polymeric micelles at various drug loadings ratio over time (polymer/drug w/w ratios of 10/1, 10/2, 10/4 and 10/8). Although some T3 micelles also show the sphere-to-worm transition, this transition is predominantly absent in this sample set.

**SANS data on T2 micelle systems**

Attempts were made to fit the blank micelle SANS data using a uniform scattering length density cylindrical model. However, the results showed poor agreement with the data at higher scattering angles (not shown). Calculated contour length and cylinder radius were also inconsistent with the light scattering and cryo-TEM results. A core-shell cylindrical model provided poor agreement with the scattering profiles for both blank and lower drug concentration solutions, where a worm-like micelle structure was plausible. Instead, small angle neutron scattering data were fitted using the following functional form:

$I\left( q \right)=\frac{A}{q^{n}}+I_{micelle}+B$,

where the first term describes Porod scattering from clusters of spherical micelles, B is the incoherent background and *I*_micelle_ describes scattering from individual T2 spherical micelles (blank or drug loaded) and *q* is the momentum transfer, defined as:

$$q= \frac{4\pi}{\lambda}\sin\theta$$

The micelles were modelled as core-shell spherical particles with constant ratio between the core radius and the total radius. Micelle interactions were modelled as hard-sphere interactions. The fitting parameters are shown in **Table S2**. The reduced Chi-squared values are defined as:

$$Reduced {Chi}^{2}=\sqrt{{\chi^{2}}/{(N-f-1)}},$$

where *N* is the number of data points used in the fit, and *f* is the number of degrees of freedom. For the lower scattering angle data, with large size aggregates and micelles clusters with dimensions outside the resolution of the collected data, the polydispersity significantly reduces the quality of the fits. For higher concentrations of paclitaxel and etoposide, the fitting range was limited to the higher q-range, where the main scattering contributions are due to the single micelles in solution.

**Table S2.** Fitting parameters of SANS data corresponding to the T2 solution at various drug loading ratios (polymer/drug weight ratios of 10/1, 10/2, and 10/4). Errors shown correspond to one standard deviation of uncertainty. Parameters without reported errors were kept fixed during the fitting of the SANS data.

(a)

| **Fit parameters** |  | **Olaparib** | | | **Etoposide** | | |
| --- | --- | --- | --- | --- | --- | --- | --- |
|  | **10/0** | **10/1** | **10/2** | **10/4** | **10/1** | **10/2** | **10/4** |
| **Coefficient A (x10^-6^)** | 8 | 20 | 28 | 34 | 29 | 35 | 40 |
| **Power (n)** | 3 | 2.88 | 2.82 | 2.1 | 2.7 | 2.55 | 2.0 |
| **Volume fraction** | 0.009 | 0.011 | 0.012 | 0.014 | 0.011 | 0.012 | 0.014 |
| **Core radius (Å)** | 41.3 ± 0.8 | 43.2 ± 0.6 | 45.2 ± 0.5 | 47.8 ± 0.1 | 43.0 ± 0.8 | 43.7 ± 0.1 | 48.7 ± 0.1 |
| **Overall polydispersity** | 0.36 | 0.36 | 0.34 | 0.20 | 0.40 | 0.25 | 0.20 |
| **SLD Core (x10^-6^ Å^-2^)** | 1.2 ± 0.1 | 0.7 ± 0.1 | 1.0 ± 0.1 | 0.4 ± 0.1 | 1.1 ± 0.1 | 0.9 ± 0.3 | 1.1 ± 0.3 |
| **Shell Thickness (Å)** | 29.7 ± 0.7 | 28.8 ± 0.3 | 28.8 ± 0.3 | 23.2 ± 0.4 | 26.1 ± 0.7 | 25.2 ± 0.1 | 25.4 ± 0.1 |
| **SLD Shell (x10^-6^ Å^-2^)** | 4.1 ± 0.1 | 4.5 ± 0.1 | 4.7 ± 0.1 | 6.1 ± 0.1 | 5.1 ± 0.1 | 5.4 ± 0.1 | 5.9 ± 0.2 |
| **SLD solvent (x10^-6^ Å^-2^)** | 6.4 | 6.4 | 6.4 | 6.4 | 6.4 | 6.4 | 6.4 |
| **Background (x10^-3^ cm^-1^)** | 1.8 | 0.8 | 1.0 | 1.5 | 2.0 | 0.3 | 0.2 |
| **Reduced Chi^2^** | 3.3 | 2.9 | 4.5 | 4.2 | 2.5 | 5.4 | 4.1 |

(b)

| **Fit parameters** |  | **Paclitaxel** | | | **Vismodegib** | | |
| --- | --- | --- | --- | --- | --- | --- | --- |
|  | **10/0** | **10/1** | **10/2** | **10/4** | **10/1** | **10/2** | **10/4** |
| **Coefficient A (x10^-6^)** | 8 | 210 | 400 | 800 | 250 | 550 | 600 |
| **Power (n)** | 3 | 2 | 1 | 1 | 1.6 | 1.6 | 1 |
| **Volume fraction** | 0.009 | 0.011 | 0.012 | 0.014 | 0.011 | 0.012 | 0.014 |
| **Core radius (Å)** | 41.3 ± 0.8 | 41.0 ± 0.1 | 47.6 ± 0.1 | 55.0 ± 0.1 | 46.6 ± 0.1 | 49.3 ± 0.1 | 61.4 ± 0.1 |
| **Overall polydispersity** | 0.36 | 0.20 | 0.16 | 0.15 | 0.14 | 0.18 | 0.13 |
| **SLD Core (x10^-6^ Å^-2^)** | 1.2 ± 0.1 | 0.7 ± 0.1 | 0.7 ± 0.1 | 0.7 ± 0.1 | 0.8 ± 0.1 | 0.1 ± 0.1 | 0.5 ± 0.3 |
| **Shell Thickness (Å)** | 29.7 ± 0.7 | 22.0 ± 0.2 | 21.8 ± 0.2 | 21.7 ± 0.2 | 23.5 ± 0.3 | 23.8 ± 0.1 | 27.0 ± 0.1 |
| **SLD Shell (x10^-6^ Å^-2^)** | 4.1 ± 0.1 | 5.2 ± 0.1 | 5.6 ± 0.1 | 5.4 ± 0.1 | 5.7 ± 0.1 | 5.6 ± 0.1 | 5.7 ± 0.1 |
| **SLD solvent (x10^-6^ Å^-2^)** | 6.4 | 6.4 | 6.4 | 6.4 | 6.4 | 6.4 | 6.4 |
| **Background (x10^-3^ cm^-1^)** | 1.8 | 0.0 | 0.1 | 0.0 | 0.1 | 0.5 | 0.1 |
| **Reduced Chi^2^** | 3.3 | 210 | 400 | 800 | 250 | 550 | 600 |

**Measuring effective value of critical micelle concentration (CMC) by DLS count rate of polymeric micelles**

Previous work indicates that drugs in POx micelle could have specific interaction with the backbone structure of the poly(2-oxazoline) and with components of the hydrophilic shell-forming block, dependent on loading amount. These interactions could offer additional stability to the spherical morphology, and could lower the CMC and chemical potential inhibiting the transition to worm-like structure both in terms of its thermodynamic favorability and the kinetics due to an increase of the potential barriers between morphological forms. We use here effective CMC* terminology to acknowledge that in some cases the formed aggregates (e.g., spherical micelles in the absence of any drug) are metastable and transform to worm-like micelles over time. However, since the transition is very slow (days) as a first approximation an equilibrium between the micellar aggregates and single block copolymer chains can be characterized by effective CMC*. As pointed out before the effective CMC* can be considered as a surrogate metric of the thermodynamic stability of polymeric micelles ^[1]^. One additional consideration of the effective CMC* in this study is that the drug loaded micelles undergo partitioning of the solubilized drug into aqueous phase upon dilution ^[2]^. This can complicate the interpretation of the CMC* value especially at high drug loadings since the drug loaded micelles are *de facto* a multicomponent system comprising both the drug and the block copolymer components. We have selected DLS as the method for estimating the effective CMC* values as point of deflection (e.g., slope change) in the intensity of light scattering indicating disassembly of the block copolymer or block copolymer and drug aggregates. With this consideration in mind, we characterized the changes in the CMC* and used this as a metric for micelle stability at various drug loadings. Overall, as drug loading increased, the CMC* value gradually decreased indicating an increase in stability against dilution - likely the result of increased drug-polymer intermolecular interactions in the micelle. While all the drugs studied displayed the same trend, they did so with different magnitudes. For instance, vismodegib and paclitaxel, which belong to the micelle elongation-inhibiting group, produce greater shifts in the CMC* when encapsulated into micelles suggesting greater micelle stabilization than etoposide or olaparib. It is clear that the lower CMC* and the increased stability of the drug-loaded micelle (estimated as 82.2 $\mu$g/mL and 53.3 $\mu$g/mL for vismodegib and paclitaxel and 198.3 $\mu$g/mL and 175 $\mu$g/mL for etoposide and olaparib at a 10/4 polymer/drug w/w ratio) explain why the elongation behavior was different across the different drugs.

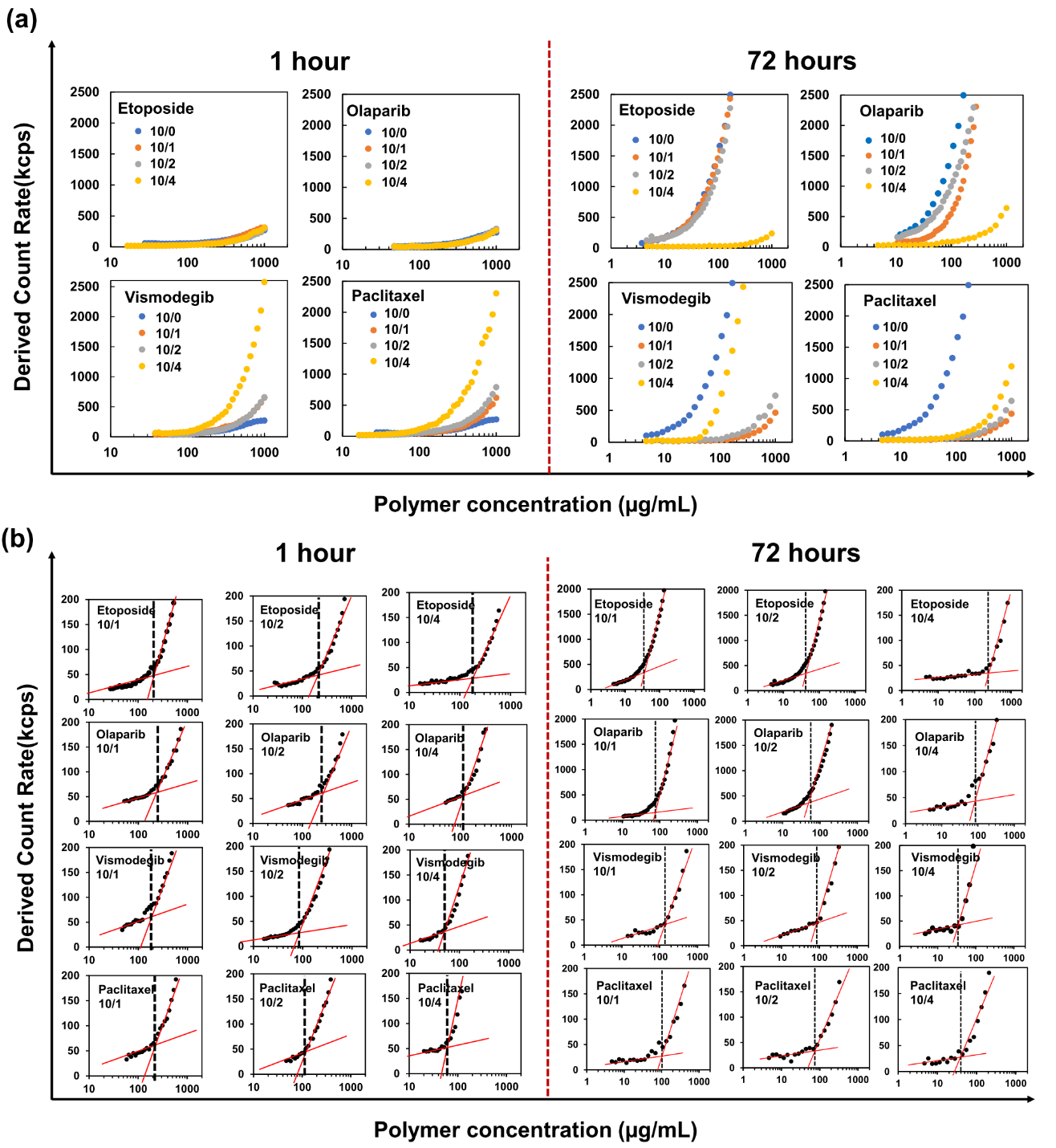

**Figure S6.** **(a)** Derived count rate of polymeric micelles with gradual dilution of samples at various drug loading ratios (polymer/drug w/w ratios of 10/0, 10/1, 10/2, and 10/4). **(b)** Derived count rate (kilo-counts per second (kcps)) plot with enlarged Y-axis used to measure the effective CMC*; the point of slope change in the derived count rate was defined as the CMC*. A T2 concentration of 1 mg/mL was used in distilled water at 25^o^C.

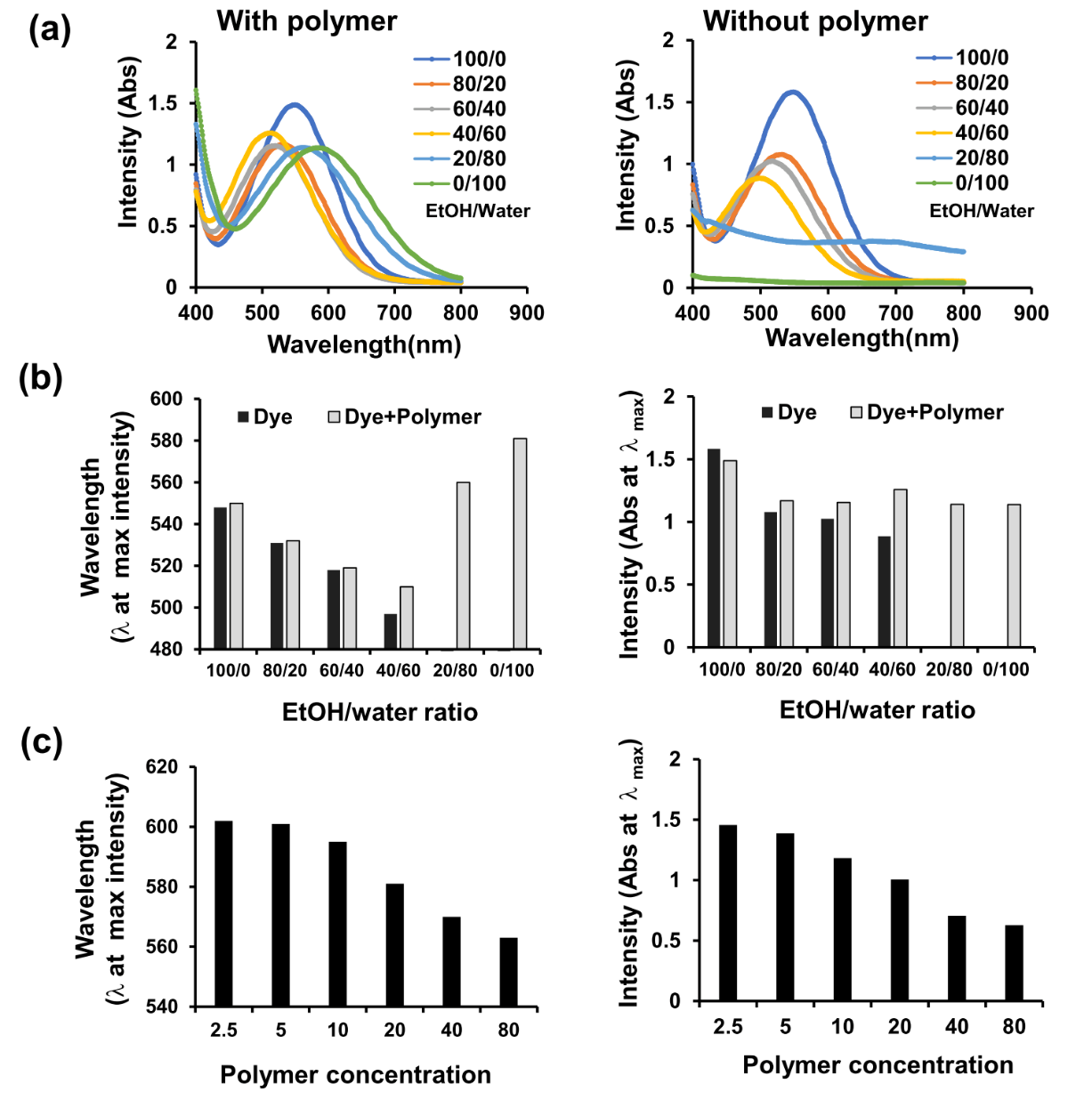

**Figure S7.** **(a)** UV-Vis absorption spectra of Reichardt’s dye (RD) at various ethanol/water solvent volume ratios (100/0, 80/20, 60/40, 40/60, 20/80, and 0/100), with and without T2 (10 mg/mL). **(b, c)** Wavelength at the maximal absorbance (λ_max_) and maximal absorbance (Abs) of RD in **(b)** ethanol/water mixtures (corresponding to spectra in plots (**a**)) and in **(c)** aqueous solutions at various polymer concentrations. RD (0.5 mg/mL, 25 ^o^C) was diluted either in **(a, b)** distilled water or **(c)** saline (0.9 % NaCl). The λ_max_ of RD decreases as the polarity of the microenvironment increases; from a λ_max_ of 548 nm in ethanol (100 %) to a λ_max_ of 497 nm in the ethanol/water mixture (40 % / 60 %). The λ_max_ of the RD in the POx micelles was higher compared to both the pure ethanol and the ethanol-water mixture, at any polymer concentration above the CMC. This is due to the relative hydrophobic environment maintained by 2-butyl-2-oxazoline core. However, as the polymer concentration increased, λ_max_ decreased, suggesting transfer of the dye within the micelles in polar environments. This was accompanied by a decrease in the dye absorbance, indicating specific interactions between the dye and the polymer.

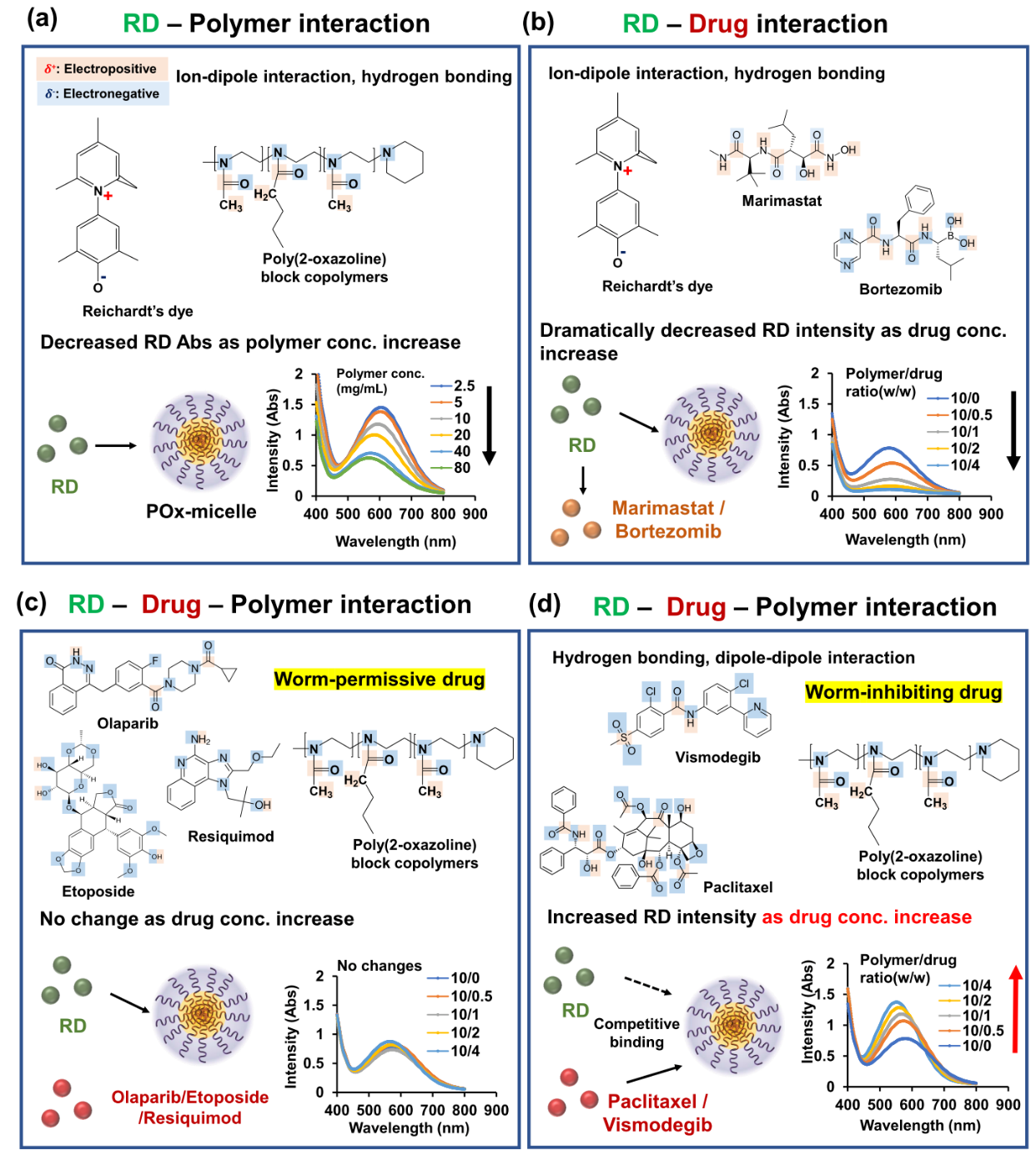

**Figure S8.** Schematic representation of the RD, polymer, and drug interactions resulting in the observed changes in dye absorbance intensity. **(a)** The betaine RD molecule may interact with the repeating amide bond motifs in the POx block copolymer resulting in attenuation of the dye absorbance. **(b)** Co-loading of the RD in the micelles with marimastat and bortezomib having a repeating amide bond motif results in further attenuation of the dye absorbance intensity. **(c)** The drugs without the amide bond motifs do not interact with the RD and therefore do not change its absorbance intensity. **(d)** Worm-inhibiting drugs such as paclitaxel and vismodegib interact strongly with the same binding sites in the polymer and displace the RD, increasing the dye absorbance intensity.

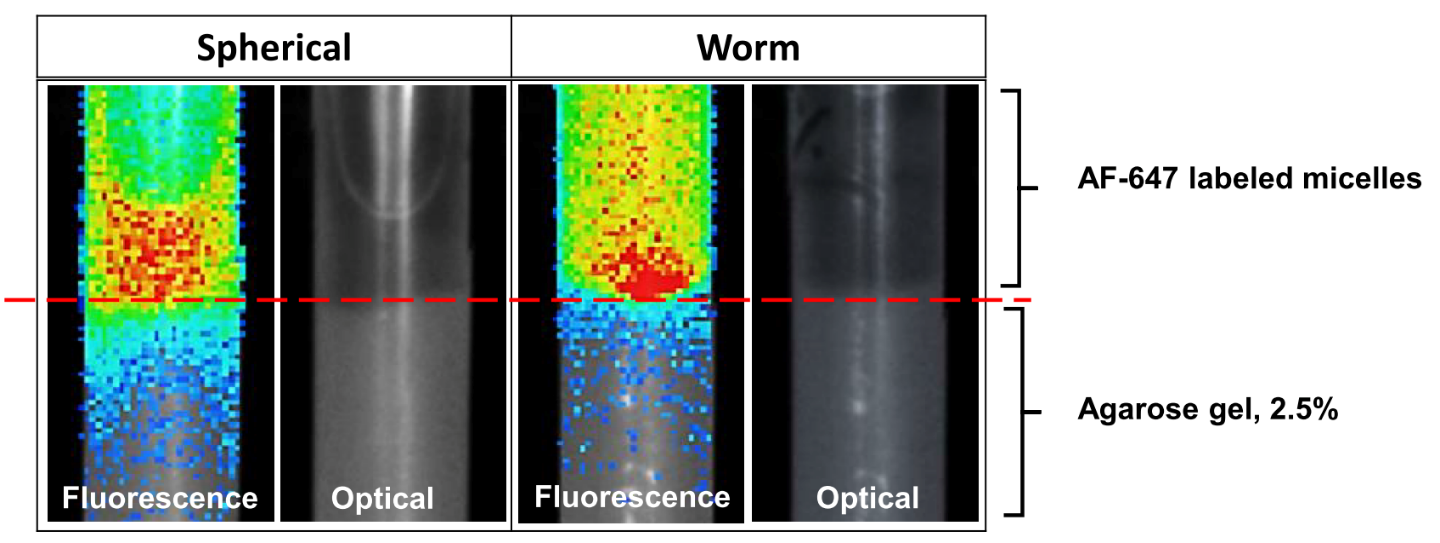

**Figure S9.** Simulation of drug-loaded micelle penetration. Dye-labeled spherical or worm-structured micelles (10/1 polymer/drug ratio, 10 mg/mL of T2 in 0.9 % NaCl) were added into a column containing 2.5 % agarose gel and incubated for 24 hours under a water flow. The red dashed line indicates the border line between the water and agarose gel. Spherical micelles showed better penetration in the model.

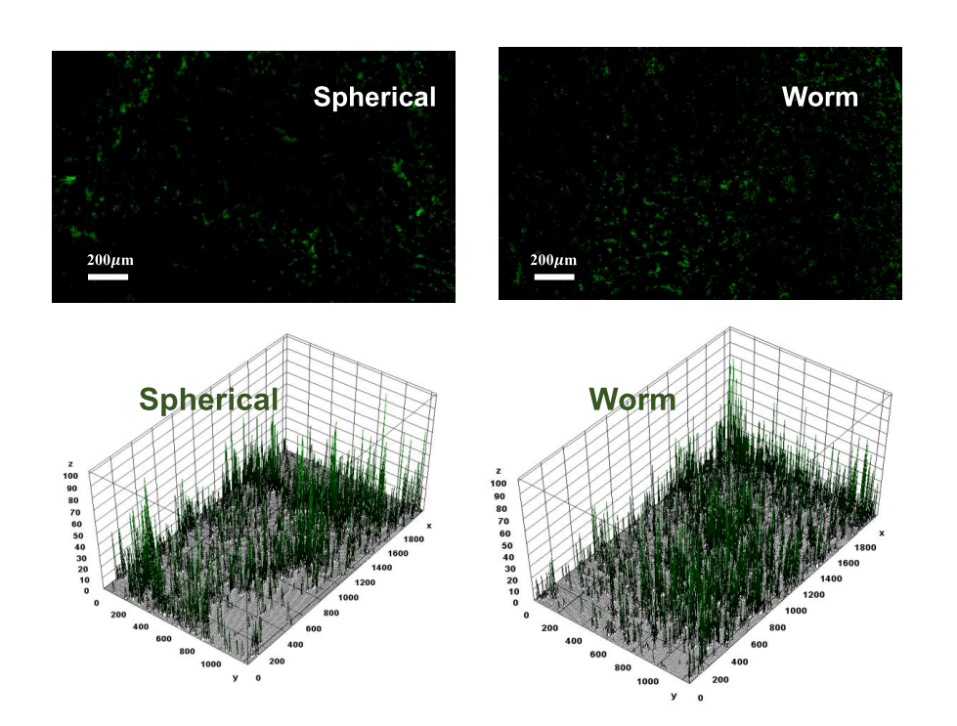

**Figure S10.** Blood vessels were stained with an antibody against CD31. CD31 fluorescence intensity at tumor sections was visualized using 3D surface plots (Image J). X and Y axes represent the area of the tumor sections, and the Z axis represents the CD31 (Green) intensities. Polymer distributions in the same tissue sections and respective 3D surface plots are shown in **Figure 9c**.

**Table S3.** AUC and clearance of polymer and drug in tumor and plasma after a single injection of olaparib (10 mg/kg) in the spherical (S) and worm-like (W) micelles (10/1 composition).

| **Location** | **PK Parameter ^a^** | | **Polymer** | | | **Drug** | | |
| --- | --- | --- | --- | --- | --- | --- | --- | --- |
|  |  |  | **S** | **W** | **p^b^** | **S** | **W** | **p^b^** |
| Tumor | AUC | Very early (0-1 h) | 45.1 | 29 | - | 2.9 | 1.7 | - |
|  |  | Early (0-2 h) | 95.5 | 66.2 | - | 4.6 | 2.9 | - |
|  |  | Late (2-24 h) | 1371 | 1573 | - | 9.3 | 8.5 | - |
|  |  | Very late (8-24 h) | 1016 | 1230 | - | 5.7 | 5.3 | - |
|  |  | AUC_all_ | 1466 | 1639 | ns | 13.8 | 11.4 | * |
| Plasma | AUC | Very early (0-1 h) | 1146 | 1248 | - | 3.3 | 3.8 | - |
|  |  | Early (0-2 h) | 1830 | 2175 | - | 5 | 5.8 | - |
|  |  | Late (2-24 h) | 6169 | 7622 | - | 9.7 | 8.8 | - |
|  |  | Very late (8-24 h) | 3314 | 4035 | - | 5 | 3.7 | - |
|  |  | AUC_all_ | 8000 | 9796 | * | 14.7 | 14.6 | ns |
|  | CL (mL/h) | | 0.39 | 0.29 | * | 17.9 | 12.0 | ns |
|  | CL2 (mL/h) | | 1.92 | 0.53 | ns | 34.8 | 24.0 | ns |

^a^ AUC_all_ – Area under the curve from the time of dosing to the time of the last observation; CL- clearance blood; CL2 blood to organ intercompartmental clearance for the two-compartment model shown in **Figure S11**; ^b^ Significance S vs. W groups: *p < 0.05.

To obtain an in-depth view of the PK and tumor distribution of the drug and the polymer, we analyzed four different AUC periods in addition to the AUC_all_ (**Table S3**): “very early” (0-1 hour), “early” (0-2 hour), “late” (2-24 hour), and very late (8-24 hour). Based on this analysis, we concluded that the drug and the polymer in the spherical micelles exhibited higher tumor AUC compared to the worm-like micelles during the early periods. The worm-like micelles also exhibited significantly decreased blood clearance (Cl) compared to the spherical micelles, as well as decreased intercompartmental clearance (Cl2) for both the polymer and the drug.

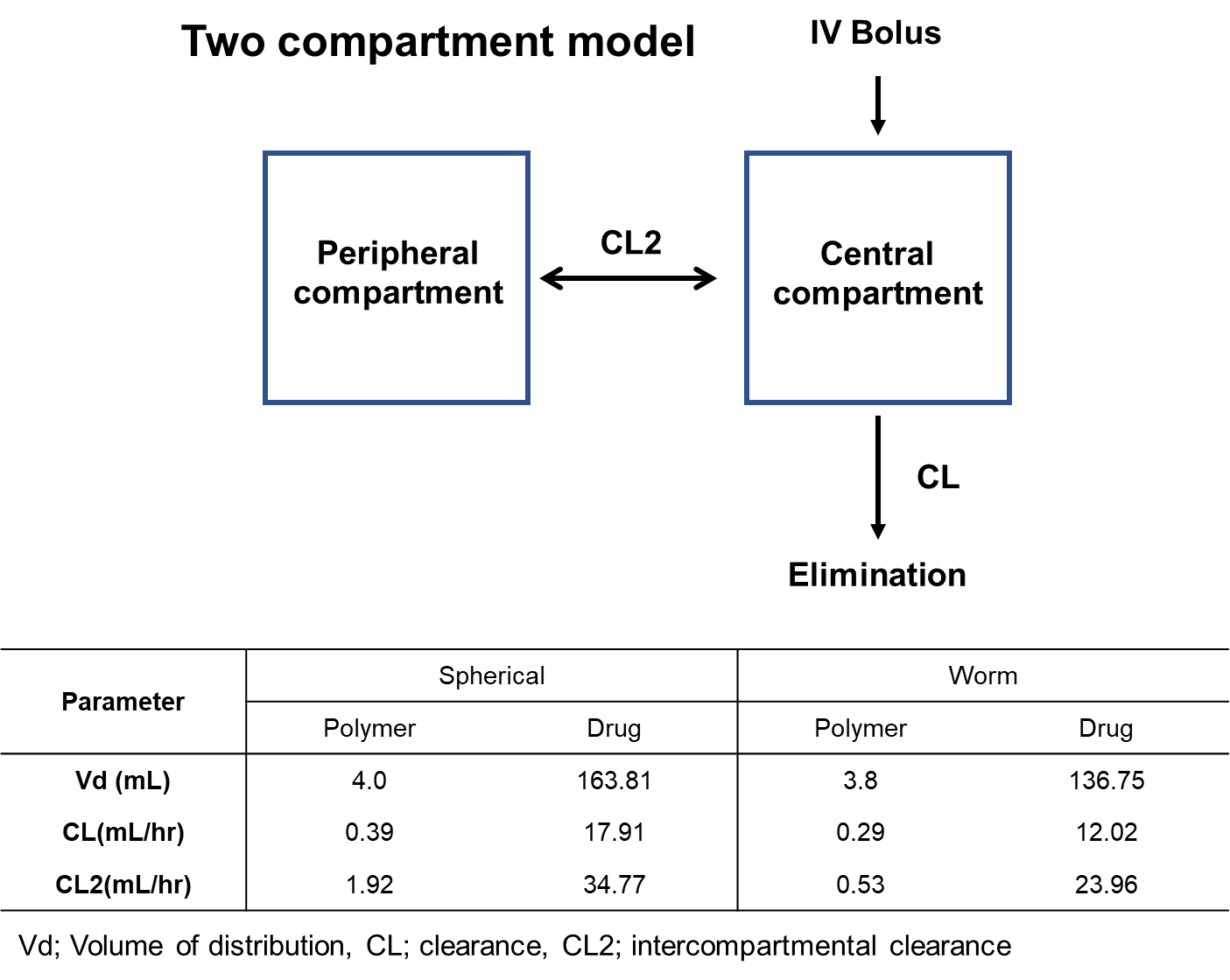

**Figure S11.** Two-compartment model describing the delivery of the polymeric micelles to a tumor. The drug is administered as a bolus in the form of spherical or worm-like micelles and is subsequently distributed between the central compartment (plasma) and peripheral compartment (including tumor). The two-compartment model with IV-bolus dosing was the best fit for polymer and drug in both morphologies. Notably, the olaparib intercompartmental clearance (Cl2) was higher for the spherical micelles than the worm-like micelles, though not with statistical significance. This indicates that the drug in the spherical micelles is able to travel faster to tissues throughout the body (e.g., tumor), accumulating faster. The same phenomenon is seen for the polymer in spherical micelles. The intercompartmental transfer in these micelles is 4 times higher than that in the worm-like micelles. The rapid transfer of both the drug and polymer in the spherical micelles (i.e., to additional tissues) suggests that they can also rapidly accumulate in other tissues.

**Disclaimer**

Certain commercial equipment, instruments, suppliers are identified to foster understanding. This does not imply recommendation or endorsement by the National Institute of Standards and Technology, nor does it imply that the materials or equipment identified are necessarily the best available for the purpose.
